# Quantifying Conformational Heterogeneity of 3D Genome Organization in Fruit Fly

**DOI:** 10.1101/2025.05.24.655945

**Authors:** Samira Mali, Igor S Tolokh, Erik Cross, Alexey V Onufriev

## Abstract

The three-dimensional (3D) organization of interphase chromatin in eukaryotes is complex; details of the corresponding genome structures vary stochastically from cell to cell. Here, we propose a metric to quantify the cell-to-cell heterogeneity of the 3D chromatin conformations in ensembles of single cells: *Conformational Heterogeneity* (*C.H.*) is defined as the standard deviation of the ensemble distribution of the per cell average Euclidean inter-loci distances ⟨*R_s_*⟩, for a given genomic separation *s* between the loci. We have used the metric to examine and quantify in detail the cell-to-cell heterogeneity of conformations of the interphase X chromosome in fruit fly generated via three distinctly different modeling approaches, which take experimental Hi-C data as input. Two of the approaches use bulk Hi-C and lamina-DamID data, while the third relies on single-cell Hi-C maps. An algorithm is proposed to facilitate comparison of *C.H.* of models constructed at different resolutions, and to examine the behavior of conformational heterogeneity with increasing model resolution. Higher resolution models show a greater *C.H.*, in general. The impact of the model resolution is strongest near the genomic distance *s* corresponding to the resolution limit of the model, and diminishes for larger genomic distances: extrapolating the resolution from approximately 14 kb to 2 kb has little effect on the *C.H.* beyond ∼100 kb. All chromatin models examined in this work show a very similar trend of monotonically increasing structural heterogeneity with *s*, up to the genomic TAD size; beyond that, significant differences arise, with the model based on single-cell Hi-C showing nearly opposite trend compared to the two models that use bulk Hi-C data. We attribute these major differences to relatively subtle differences in the modeling approaches, which we discuss. Based on the analysis, we propose to explore the possibility of inclusion of bulk Hi-C data into training of chromatin models that are based on necessarily limited single-cell Hi-C data. Within our computational approach, depletion of nuclear lamins leads to increased structural heterogeneity at nearly all genomic separations, with the potential implication that cell functions that depend on chromatin structure might be more variable within lamins depleted nuclei compared to the wild type.

## Introduction

The three-dimensional (3D) organization of chromatin [1] within a cell nucleus plays a pivotal role in the regulation of gene expression and cell repair [2–9]. An important consideration is the naturally occurring heterogeneity/variability [10, 11] of chromatin structure from cell to cell. This heterogeneity of the 3D chromatin organization includes different mutual spatial arrangements of the chromosomes [12, 13], which varies from nucleus to nucleus, as well as variations in chromatin packing on smaller scales within each chromosome, resulting in inherent heterogeneity of chromatin structures of the same cell type in the tissue.

To investigate the chromatin organization, chromosome conformation capture (3C) technique [14], and various 3C-derived methods – 4C [15], 5C [16] and Hi-C [17] – have been developed, among others. In particular, the Hi-C technique provides a matrix of contact frequencies between pairs of genomic loci, with the current genomic resolution up to 1 kb [18]. These methods have revealed a wealth of information about how chromatin folds in the tiny space of the nucleus [19], including the existence of topologically associating domains (TADs) and higher-order chromatin structures such as A/B compartments and chromosome territories [20–22]. However, several challenges exist in interpretation of the Hi-C data: one important limitation is that the standard bulk (population averaged) Hi-C method yields a single contact frequency map representing an average over a very large ensemble (millions) of individual nuclei, which helps reduce the noise in the underlying experimental data, but at the cost of information loss.

To complement population-based chromosome conformation capture methods, the single-cell Hi-C (scHi-C) technique [23–26] provides chromatin contact maps of individual nuclei. The approach helped to confirm the existence of Topologically Associating Domains (TADs) and compartments in individual cells [27, 28] and offers insight into cell-to-cell heterogeneity of chromatin organization within the nucleus, chromosome compartment dynamics and evolution during cell cycle [29]. The availability of scHi-C data also made it very clear that chromatin conformation features of individual cells are not identical to population-average chromatin folding patterns [23, 26, 28–30]. However, technical limitations of the scHi-C technique limit the amount of information that can be inferred from these maps at present – the contact frequency maps reported in a single study are very sparse and few. For example, a recent scHi-C study [28] of chromatin in fruit flies yielded only about 88 scHi-C maps, of which only 20 were used for further analysis by the authors, as these 20 did not have PCR duplicates and showed a rapid rise in unique contacts with sequencing depth. These limitations contribute to the lack of quantitative understanding of exactly how heterogeneous/diverse chromatin structures within the same cell type are.

Moreover, even perfect Hi-C maps, bulk or single-cell, do not provide direct structure of chromatin in the 3D space. Attempts to infer a single chromatin conformation from a single Hi-C map may lead to geometric inconsistencies, violating triangle inequality [31] – a fundamental feature of Euclidean space. The issue is further complicated by the fact that chromatin conformations are not static, evolving in time [13, 32–39] even within the same phase of the cell cycle, further contributing to natural conformational heterogeneity of chromatin in living cells.

In contrast, microscopy, including fluorescence in situ hybridization (FISH), can access distances between loci in Euclidean space, thus providing direct measurements of 3D chromatin structures in individual cells. However, the resulting “distance matrices” are very sparse, with the current limit of several thousand loci [40].

Computational models of chromatin organization in 3D space [13, 37, 41–47] are therefore essential to complement the experiment in this area, helping to make connections between chromatin structure and its biological function [48, 49]. These models [13, 41, 50–54] often use the Hi-C data for parameterization and validation, including bulk [13, 41] or, more recently, single-cell [28] Hi-C data. Given that each type of Hi-C map has its own limitations, exploring similarities and differences between the resulting 3D structures is of value. However, a meaningful comparison between ensembles of chromatin structures is non-trivial, even for the same organism and cell cycle stage: the same bulk Hi-C map can result from different distributions of scHi-C data. And, as mentioned above, a 3D structure of the genome is more than its Hi-C map. As a useful step towards developing a computationally facile, yet intuitive and informative approach to comparison of complex chromatin structures in 3D space, here we propose a strategy to compare structural heterogeneity of 3D chromatin structures of fruit fly – a well studied organism with only four chromosomes, for which several distinct models of 3D genome organization at 10-100 kb resolution are available.

We begin by proposing and exploring a metric designed to characterize structural heterogeneity of 3D genome organization along the genome by a single number, appropriate for each genomic scale of interest. To our knowledge, although several studies have discussed the conformational heterogeneity of chromatin in various contexts [10, 11], none has proposed a single, computationally facile metric for it that may serve as an analogue of the first moment of a distribution. Also note that we seek to quantify the heterogeneity of an ensemble of polymer conformations in 3D, not its population-averaged Hi-C map [55]. Here we will fill this gap, and exemplify the use of the new metric on three distinct models of 3D organization in fruit fly. An algorithm is developed and applied to facilitate comparison between models available at different spatial resolutions. We will also explore how chromatin conformational heterogeneity depends on the model resolution, including the theoretical limit of a single base-pair resolution. To demonstrate how the new metric can be used in practice, we explore the effect of nuclear lamins depletion on cell-to-cell structural heterogeneity of fruit fly chromatin.

## Methods

We begin this section by briefly introducing three computational approaches previously used to model 3D genome organization of *Drosophila* nuclei: the resulting published structural models are employed in this study. We then describe in detail how we pre-process the available data to select the desired conformation ensembles. Then, we introduce our MC-TAD algorithm, which reconstructs, to a specified resolution, the chromatin conformations inside a TAD. Next, we describe how the MC-TAD is used to “up-convert” chromatin conformations of lower resolution models to facilitate comparisons between different models. Details of these procedures are presented in the Supporting Information (the S1 Text). The S1 Text also contains multiple additional auxiliary details such as unit conversions.

### Three chromatin modeling approaches

We analyze three sets of *Drosophila* X chromosome conformations resulting from three different computational approaches to model the spatial organization of chromatin: the Li et al., 2017 [41], Ulianov et al., 2021 [28] and Tolokh et al., 2023 [13] chromatin modeling approaches. Relatively large (∼20 Mb) heterochromatin regions of the X chromosome, not considered in Ulianov et al., 2021 models, have been excluded from the Li et al., 2017 and Tolokh et al., 2023 models to facilitate direct comparison. Thus, only euchromatin portions of the X chromosome are analyzed in this work. To reconstruct the 3D structures of chromosomes, these three works use completely different computational approaches: simulation of chromosome dynamics using Molecular Dynamics (MD), relaxation of a polymer with additional bonds using Dissipative Particle Dynamics (DPD) [56] and Machine Learning-like methods for generating and selecting chromatin structures. We chose these three approaches because they are relatively recent works that generated a large number of 3D conformations of single X chromosome [28] or entire *Drosophila* genome [13, 41]. Each chromatin conformation can be considered as a snapshot of a single nucleus, which we refer to as *single cell*.

A key distinctive feature of the Ulianov et al., 2021 [28] approach, most relevant to our purposes, is that it aims to create a model of 3D genome organization to match a set of individual reference *single cell (sc)* Hi-C maps they constructed from their experimental data. In contrast, in both Li et al., 2017 [41] and Tolokh et al., 2023 [13] works, the reference data used to generate chromatin configurations are the TAD-TAD contact probability matrix derived from the bulk (*ensemble-averaged*) Hi-C map [20], and TAD-Nuclear Envelope (NE) contact probability data (lamina-DamID data) [57].

#### Chromatin structure models of Ulianov et al., 2021

This study [28] employs a single-cell Hi-C (scHi-C) technique conducted on 88 asynchronously growing *Drosophila* male Dm-BG3c2 (BG3) neuronal cells. Out of the initial scHi-C data, 20 single-cell libraries were selected for additional deep sequencing and generating scHi-C maps at 10-kb resolution. Using these scHi-C data and applying dissipative particle dynamics (DPD) polymer simulations [56], Ulianov et al. [28] created 20 models of *Drosophila* haploid X chromosome at 10-kb resolution. These models allowed to reconstruct chromatin folding patterns within TADs, at middle genomic distances, and at the scale of the TAD region of X chromosome.

In the simulations, a polymer chain of 2,242 equal beads with the undeformed bond length of 0.5 DPD arbitrary units (a.u.) and 29,702 solvent beads undergo DPD within the simulation box of 22 × 22 × 22 DPD a.u. Additional harmonic bonds (initially overstretched) between non-adjacent polymer beads were added according to the scHi-C contact data in each of 20 models. These bonds are retained or removed based on the results of several sets of 20k steps of equilibration for each model.

**Selection of the representative ensemble.** The final equilibrated conformations of all 20 *single cell* models of X chromosome are used for our analysis.

#### Chromatin structure models of Li et al., 2017

In this work [41], a machine learning framework was used that integrates experimental data for a comprehensive analysis of genomic structural features of chromatin conformations. Briefly, Li et al. [41] derived a coarse-grained TAD-TAD contact probability matrix (*A*) and a TAD-NE contact probability vector (*E*) from the available experimental bulk Hi-C map [20] and lamina-DamID data (chromatin-NE interaction data) [57] for *D. Melanogaster* nuclei, accordingly. The authors generated a population of chromatin structures, whose TAD-TAD and TAD-NE contact frequencies are consistent with both the matrix *A* and the vector *E*; the resulting model Hi-C map provides, by construction, a very close match to the experimental Hi-C map, having the average column-based Pearson’s correlation coefficient with the experimental map of 0.984. The resulting 3D models of fruit fly chromatin are at TAD resolution, approximately 100 kb within the embryonic *Drosophila* genome. The 10,000 selected genomic structures characterize the TAD-TAD and TAD-NE interactions across the entire diploid genome.

**Converting the two-chain models of Li et al., 2017 to a single chain representation.** Both Ulianov et al., 2021 models (male haploid X chromosome of *Drosophila* BG3 cell line [58]) and Tolokh et al., 2023 models use single polymer chain to represent haploid or paired diploid chromosomes, accordingly, while Li et al., 2017 models utilize two polymer chains representing two paired homologous female X chromosomes. To compare all three models on an equal footing, we convert the two-chains conformations of paired X chromosomes from Li et al., 2017 models to a single chain representation. Each pair of homologous TADs represented by two beads in that model, is replaced by a single bead, placed at the center of mass of the pair. The radius of each new single bead remains the same as the radius of the initial beads in the corresponding homologous pair. The resulting model of X chromosome is a chain of 184 beads of variable size representing chain of TADs. The average radius of the beads in this model of X chromosome is 0.072 microns.

**Selection of the representative ensemble.** For consistency with the number of X chromosome conformations (20) in the conformational ensemble of Ulianov et al., 2021 models, we randomly select 20 conformations from the complete set of 10,000 available chromatin structures. We repeat this data sampling procedure 10 times to obtain 10 different ensembles with 20 conformations in each. For each of these ensembles, the proposed metric (*Conformational Heterogeneity* (*C.H.*)) is calculated independently, and the corresponding error bar (standard deviation) is estimated.

Figs 7 and 8 illustrate the behavior of ⟨*R_s_*⟩ measures for only one of these 10 sets. In Fig 9b and S1 Table D in the S1 Text, the reported bulk (ensemble averaged) values include the error bars, estimated based on all the 10 sets with 20 structures in each.

#### Chromatin structure models of Tolokh et al., 2023

In this study [13], several models of embryonic *Drosophila* female interphase nuclei (with different mutual arrangements of chromosomes) were developed and simulated to generate a large set of chromatin conformations. In these models, the diploid female chromosomes, represented by chains of beads (representing the chains of TADs and heterochromatin regions), were confined to a spherical enclosure representing the NE. The interaction parameters of the models were tuned to reproduce the TAD-NE contact probability data [57] and the coarse-grained TAD-TAD contact probability matrix, derived by Li et al. [41] from the experimental bulk Hi-C map [20]. Each TAD related bead in the models represents two paired homologous TADs with approximately 100 kb of DNA in each, resulting in one polymer chain for each of two paired homologous chromosomes in the diploid female *Drosophila* genome. This homologous TADs and chromosomes pairing is observed experimentally [59–62], and explained within a theoretical model [62]. The Langevin dynamic was used to simulate 18 nuclei and generate 18 trajectories, each corresponding to approximately 11 hrs of the biological time evolution of the interphase chromatin. The relation between the experimental and simulation timescales was determined by comparing the calculated and experimental mean square displacement (MSD) of a chromosomal locus as a function of time, see Ref. [13] for details. The simulations took approximately 10 days on 18 computing cores.

The bulk Hi-C map derived from the models, that is the map calculated using model-generated trajectories, has Pearson’s correlation coefficient with the experimental Hi-C map 0.954. We would also like to note that ⟨*R_s_*(*s*)⟩ dependencies calculated for the chromatin conformations of these models are in a very good agreement with the corresponding experimentally determined dependencies for the inter-loci distances in *Drosophila* embryo nuclei [63].

Fig 1 shows the different experimentally observed topologies (mutual arrangements of the chromosomes [12]) of the *Drosophila* chromatin considered in these models.

**Fig 1.**
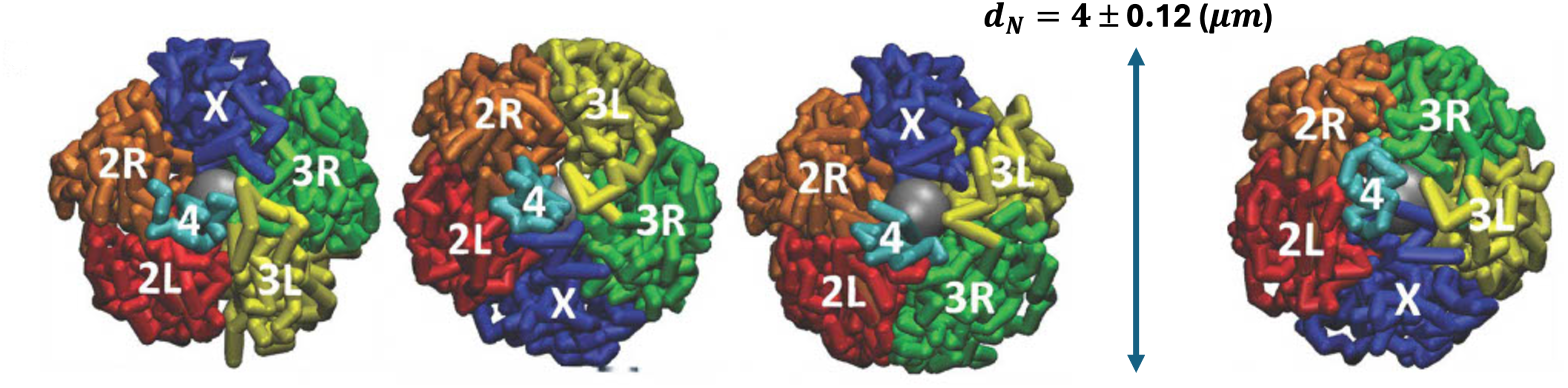
Models of *Drosophila* nuclei developed by Tolokh et al., 2023 [13]. Four possible, experimentally observed [12], different mutual arrangements (topologies) of the interface chromosomes (chromosome arms 2L, 2R, 3L, 3R, chromosomes X and 4) and three slightly different sizes (*d_N_*) of the model nuclei are taken into account. 18 model nuclei (single cell nuclei) were simulated using Langevin dynamics: the generated trajectories of the interface chromatin time evolution correspond to approximately 11 hours of biological time. The movie https://github.com/Onufriev-Lab/hi-c_model_validation/blob/main/ChrX_TRANS_X3S_1min_Droso_Interphase.mp4 illustrates the time evolution of X chromosome in one of these models.

The chain of TADs in X chromosome is represented by a chain of 184 beads of variable size; the average radius of the bead is 0.091 microns.

**Lamins depleted model nuclei.** Lamins depletion is modeled by setting to zero the affinity of TADs, that contain lamina-associated domains (LADs), to the NE (see Ref. [13] for details). The selection of representative ensemble is the same as for the wild type nuclei.

**Selection of the representative ensemble.** Each of the 18 trajectories is represented by a set of 400,000 chromatin conformations (snapshots) and corresponds to approximately 11 hrs of interphase chromatin evolution. Each trajectory is divided into 50 equal segments, each corresponding to ∼13 min. We randomly select 1 of 50 snapshots, that represent boundaries of these segments, in each of the 18 trajectories, as shown in Fig 2, resulting in 18 random *single cells* chromatin conformations. This data sampling procedure is repeated 10 times to create 10 different ensembles of conformations. For each of these 10 sets of 18 conformations, the proposed *Conformational Heterogeneity* metrics are independently calculated, allowing to estimate the error bars to assess statistical reliability of the results.

**Fig 2.**
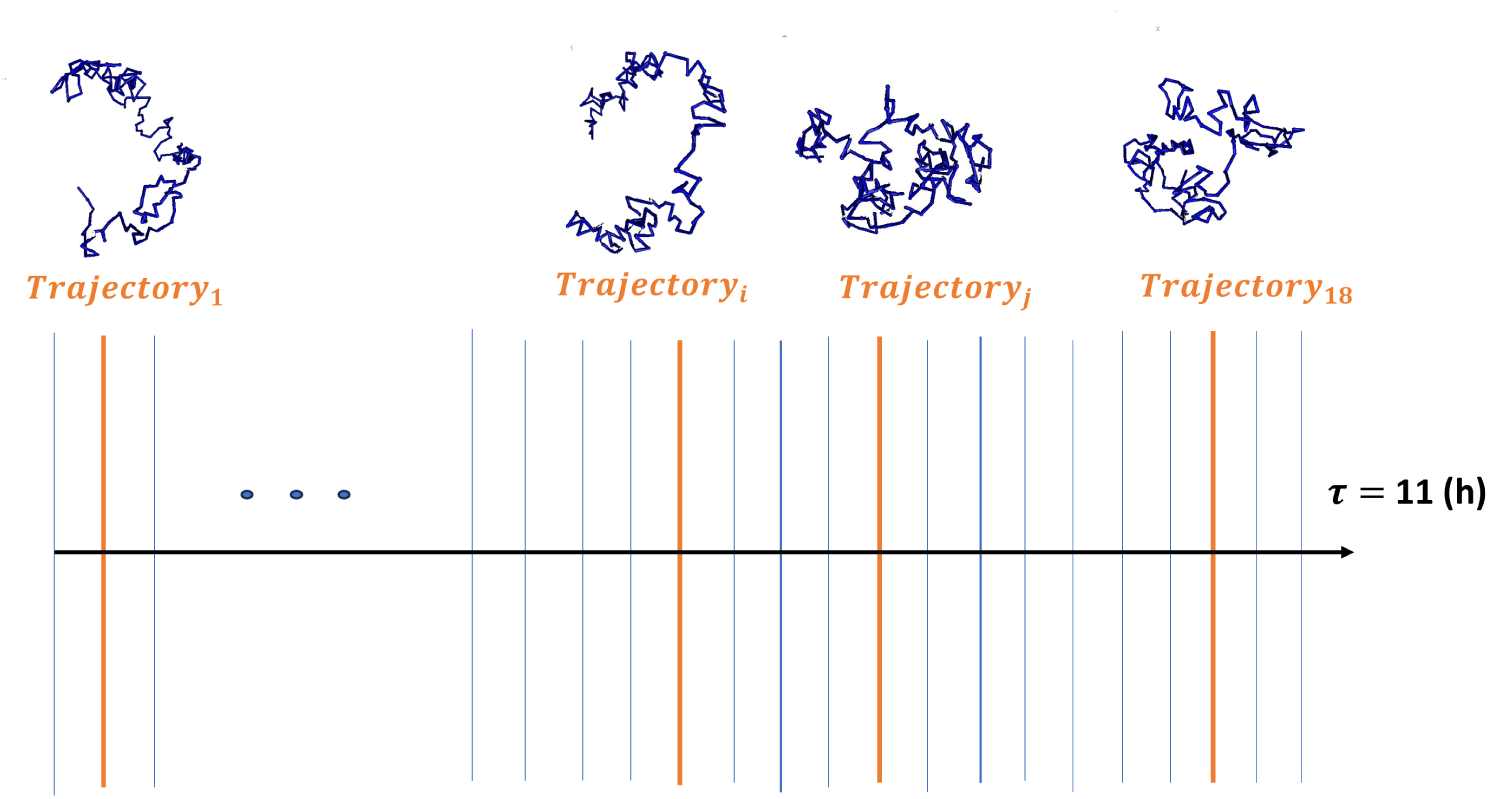
The data sampling procedure used to generate single cell conformational ensembles from the temporal sequence of conformations from Tolokh et al., 2023 *Drosophila* nuclei models. We divide each of the 18 available 11-hour trajectories into 50 equal time intervals. The vertical blue lines represent boundaries of these intervals, each corresponding to one snapshot. One of these 50 snapshots (chromatin conformations) is randomly selected from each of the 18 trajectories (one orange line for each available trajectory).

Figs 7-8 and 10 show the ⟨*R_s_*⟩ and relative ⟨*R_s_*⟩ curves for only 1 of these 10 sets, used for illustration. However, the average values and the corresponding statistical uncertainties (error bars), presented in Fig 9b, in Table 1 and in S1 Tables D and E in the S1 Text, are estimated using all 10 sets (each with 18 randomly chosen chromatin conformations).

**Table 1.** Values of the *Relative C.H.* of Tolokh et al., 2023 models extrapolated to 2-kb resolution, see Methods. The error bars represent standard deviation over all 10 values obtained as the *Relative C.H.* of different sets of selected chromatin ensembles, see Methods.

| Relative<br>Conformational Heterogeneity | 2 kb | 118 kb | 1 Mb | 10 Mb |
| --- | --- | --- | --- | --- |
| Tolokh 2023 | 0 | 0.17 ± 0.01 | 0.05 ± 0.02 | 0.19 ± 0.04 |

### Ensemble of space-filling fractal curves

We have used the Hilbert curve [64] as a model of space-filling fractal, which can in turn be considered as a relevant model of chromatin packaging [42]. Starting with the algorithm described in [65], and an implementation in numpy-hilbert-curve 1.0.1, we have used ChatGPT (model o3) to augment the algorithm to generate 18 replicas of *approximate* orientation-random Hilbert curves – the curve-to-curve variability, Fig 3, is intended to mimic cell-to-cell heterogeneity. Algorithmically, the variability is introduced by taking the canonical 3D Hilbert ordering of 8*^p^* points and, at each recursion depth *d* = 1, …, *p*, applying a fresh random cube symmetry (a random permutation of the *x, y, z* axes combined with independent sign-flips) to the order-1 8-point template before descending into its eight children. This per-level reorientation preserves the continuity of the curve, while yielding distinct space-filling paths. Here we have used *p* = 4, which results in curves of 4096 points each. The mean squared displacement *R*(*s*) for each of 18 such replicas follows the apparent power-law scaling:

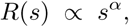

which remains close to that of the canonical Hilbert curve *α* = 1*/*3, with only small replica-to-replica fluctuations: fitting log *R_s_*versus log *s* over the interval *s* ∈ [2, 400] yields

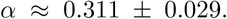

**Fig 3.**
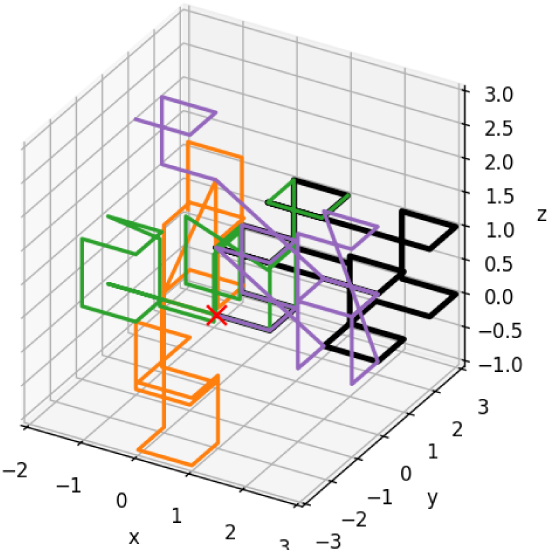
An illustration of four approximate, orientation-random Hilbert curves of 4096 points (4095 segments) each; only the first 30 points (29 segments) of each curve are shown for clarity. Due to the presence of some diagonal segments, the curves are approximate representations of the canonical Hilbert curve.

The small spread in *α* confirms that the locality and fractal scaling are preserved in our approximate orientation random Hilbert curves, while providing an additional degree of cell-to-cell heterogeneity.

To facilitate comparison with the polymer models of chromatin described above, we “coarse-grain” the genomic coordinate *s* of each *R*(*s*) curve by a factor of 2, and assign the unit segment length of each curve the genomic distance of 10 kb, which results is a fractal-like chain of 2048 “beads” that span the genomic distance from 10 kb to approximately 20 Mb, see Fig 9a.

### Ensemble of Freely Jointed Chains

Arguably the simplest meaningful “generic” representation of a polymer chain is a Freely Jointed Chain (FJC) of *s* ≫ 1 segments of length *l* each. In the case of FJC, the end-to-end distance follows normal distribution with the mean value of ⟨*Rs*⟩ = *l*√*s* and standard deviation Σ = *l*√*s*/3. Considering *s* as representing the genomic separation, for long enough chains we arrive at

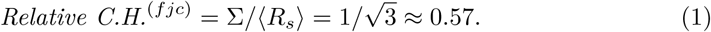

We have used ChatGPT (model o3) to generate 18 distinct random walk replicas of 2047 identical unit segments each (2048 “beads”). To facilitate comparison with the polymer models of chromatin described above, we have assigned the genomic distance of 10 kb to unit segment length of each FJC, which results in the chain that span the total genomic distance from 10 kb to approximately 20 Mb, see Fig 9a.

The mean squared displacement *R*(*s*) for each of 18 FJC replicas follows the apparent power-law scaling

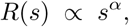

which remains close to that of the exact random walk *α* = 1*/*2, with only small replica-to-replica fluctuations: fitting log *R_s_* versus log *s* over the interval *s* ∈ [2, 400] yields

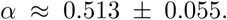

It is also reassuring that *Relative C.H.* of numerically generated ensemble of FJC of finite length, see Fig 9a is well below its upper bound (*s* → ∞) estimate provided in Eq 1.

### Calculating average distances **⟨*R_s_*⟩**

For each of the selected X chromosome conformations, we calculate average Euclidean distances ⟨*R_s_*⟩ between the centers of genomic regions separated by genomic distance *s*, as described in S1 Eq 3 in the S1 Text. To avoid complications arising from the fact that some chromatin models have beads representing TADs of significantly different size, we measure *s* in the number of beads of the same genomic size – average TAD size in X chromosome. In the case of Li et al., 2017 and Tolokh et al., 2023 models, where beads represent TADs, the average genomic size of a TAD in X chromosome is 118 kb. We use this value to convert *s* from the number of beads to genomic distance (separation) in kb in these two models. Fig 7a illustrates the relationship between ⟨*R_s_*⟩ and genomic distance *s* across different models for one set of chromatin conformations discussed previously.

### MC-TAD algorithm

In this section, we describe the algorithm “MC-TAD” (Monte-Carlo TAD), which generates all possible chromatin conformations at a specified resolution within TADs. We use this algorithm to “up-convert” the resolutions of chromatin conformations in the lower-resolution chromatin models to the conformations with higher resolution, Fig 4. The algorithm is also used to explore the effect of model resolution on structural heterogeneity of chromatin conformations.

**Fig 4.**
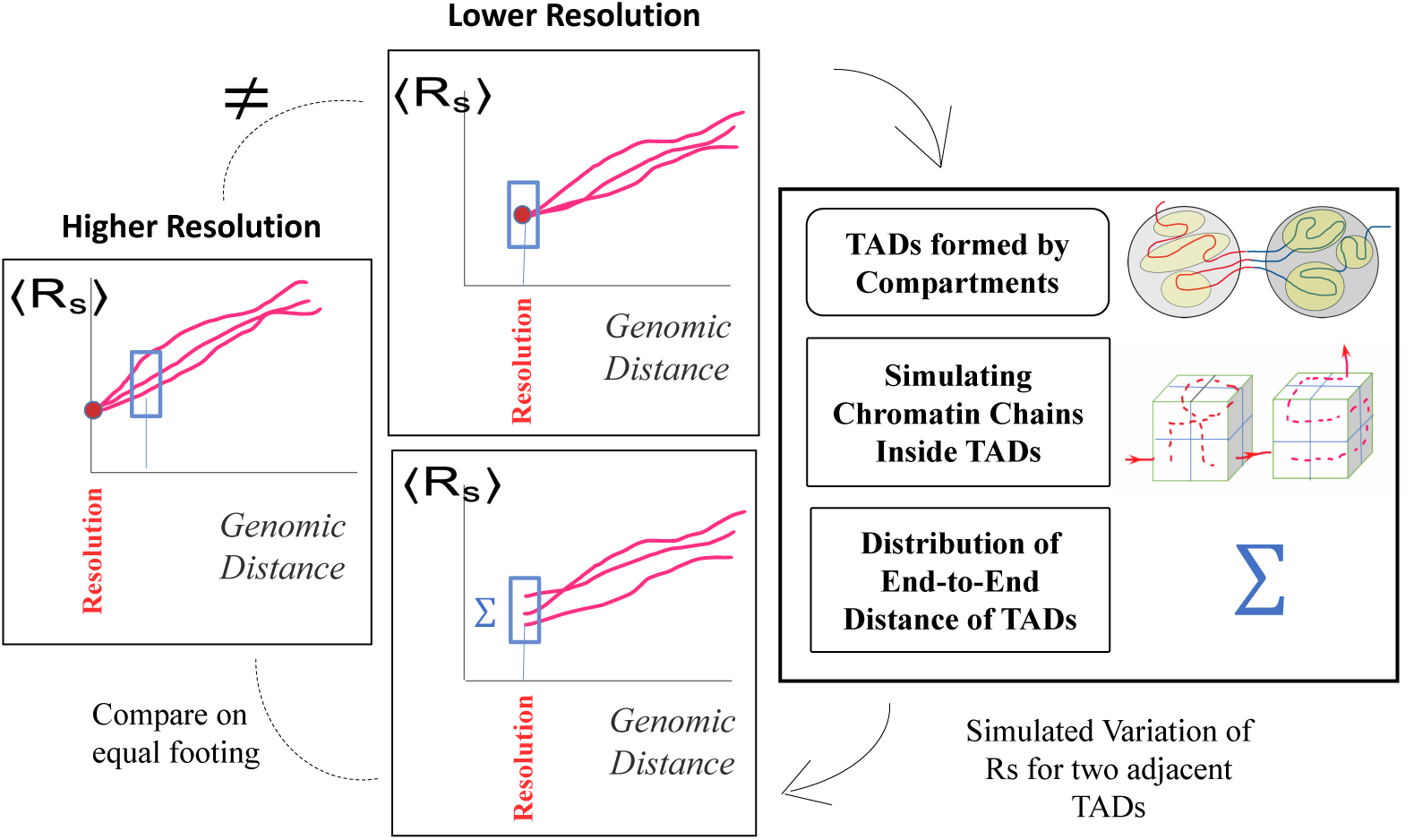
A lower resolution model of chromatin (middle panel, top) can be up-converted to match the spatial resolution of a higher resolution model (left panel) by approximating the spread of inter-loci average Euclidean distances ⟨*R_s_*(*s*)⟩ for different *single cell* chromatin conformations. The spread, Σ, is inferred from a modeled chromatin chain packing within TADs, represented by cubes which are subdivided into smaller bins to mimic the desired higher resolution (right panel). Each red curve, ⟨*R_s_*(*s*)⟩, characterizes single chromatin conformation in an individual nucleus.

#### Generating allowed chromatin conformations

Each TAD is considered as a large cube containing *N* × *N* × *N* small cubes of the same size, approximately corresponding to a bin size (resolution) in the Hi-C map [18]. In the following, we will refer to these small cubes as *bin*s. For example, one can specify *N* = 2, which results in 8 bins to model the chromatin structure of ∼118 kb *Drosophila* TAD with approximately 14-kb resolution (∼118 kb*/*8 ≈ 14 kb). We now ask how many distinct chromatin paths are possible that thread through the bins of one TAD and continue to neighboring TAD with the same resolution? These paths will represent conformations of the chromatin chain that fit inside TAD, thereby providing an approximation for the fine structure of a TAD at a given resolution. The MC-TAD algorithm generates all permissible paths, Fig 5, which are referred to as *chromatin conformations*. The total length of each path, that is the combined total length of connecting segments in a TAD, matches the genomic size of the average TAD in the *Drosophila* genome.

**Fig 5.**
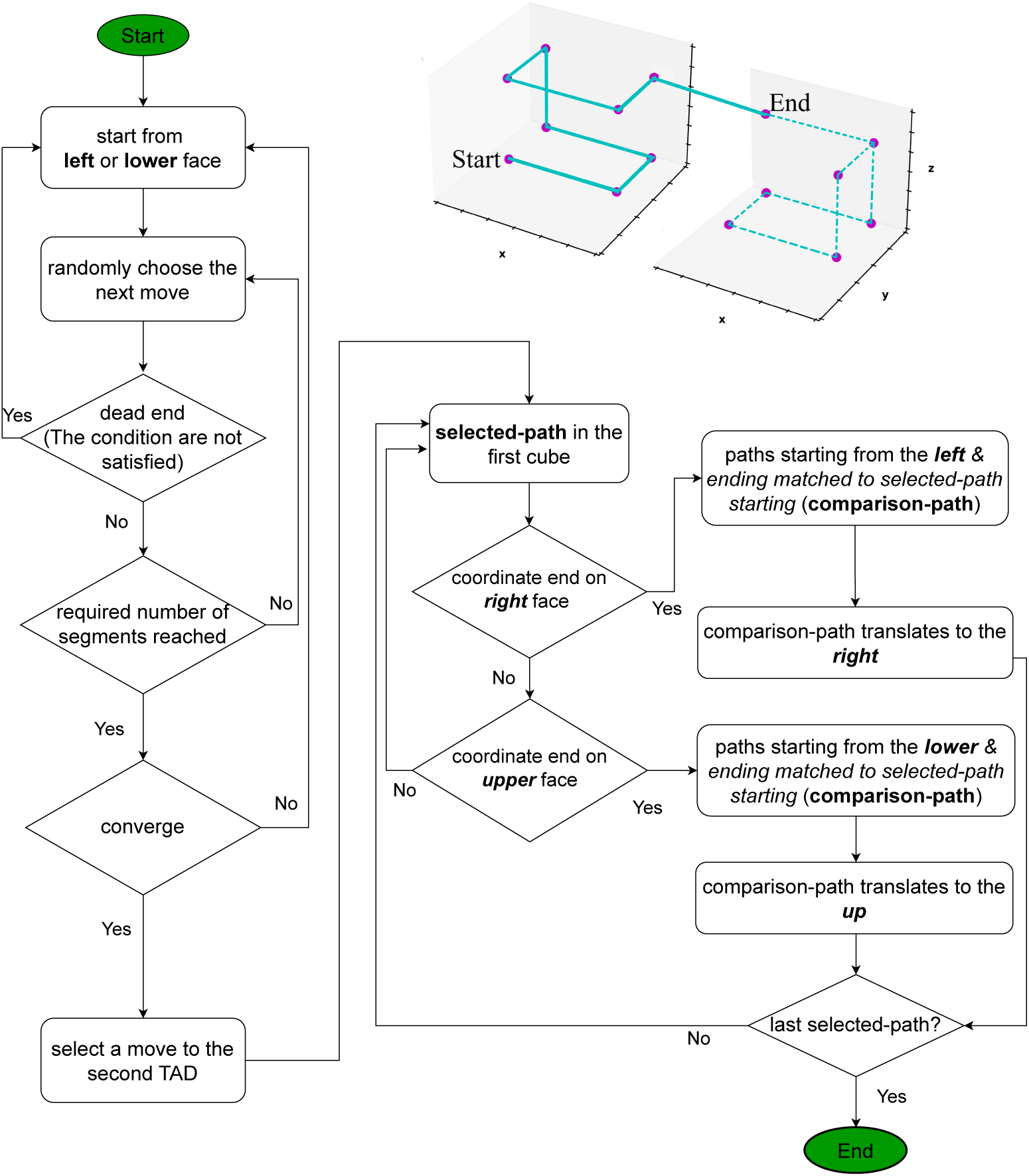
A flowchart of the MC-TAD algorithm for generating the permissible paths inside the first and second cubes representing two neighboring TADs. The generated path enters the second cube after it has traversed (*N* × *N* × *N*) − 1 full segments in the first cube. Because of a directionality of the chromatin chains between TADs and the specified path length in one TAD, a starting face in the first cube cannot coincide with the starting face in the second cube, unless the starting bin belongs to the edge of the cube. The upper right schematic illustrates one of the many permissible paths for the case of *N* = 2 (8 bins per cube) to approximate its fine structure.

Permissible paths (allowed conformations) are determined and restricted by specific biological features (see Sec. “Generating permissible conformations within a single cube” in the S1 Text) and geometrical conditions of the *chromatin conformations* that interest us. For example, while the path (conformation) must not cross over a connecting segment, i.e., the segment connecting the centers of two nearest bins along the chain, the path is permitted to cross itself at a single point, in the center of the bin, representing a self-contact. Permitting these crossings allows us to relax the condition that the path must go through each bin in the TAD, see S1 Fig A in the S1 Text for an example. This assumption speeds up the computation significantly, while having no major effect on the outcome. Once all unique, valid *chromatin conformations* within the first of the two neighboring cubes (TADs) are generated, we proceed to the second cube in the specified direction to complete the path. The next run of the stochastic algorithm generates another path; once a valid new paths are no longer generated, we declare that convergence is reached, see S1 Fig E and S1 Table A in the S1 Text for details and a convergence analysis.

Below is an example of the execution of the algorithm for the case of *N* = 2. A total of 384 unique paths are generated within the first cube. In the two neighboring cubes 98,304 unique paths are generated. A more detailed description of the MC-TAD algorithm, including consideration of higher resolution cases (*N >* 2) and convergence issues, can be found in Sec. “Generating permissible conformations within a single cube” in the S1 Text.

#### Calculating the width σ for the distribution of allowed conformations

After generating all permissible chromatin conformations, we calculate the standard deviation *σ* for the Euclidean distances *R_s_* (see Methods) for all subchains between two neighboring TADs with the contour length *s* = 118 kb. For *N* = 2, standard deviation is:

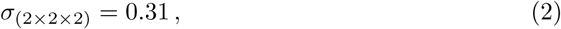

where the subscript indicates the number of bins (*N* × *N* × *N*) in each cube that represents a TAD. For simplicity, *σ* is estimated here in arbitrary units (a.u.) assuming the cube size is 1 a.u. The conversion of dimensionless *σ* values in a.u. to corresponding Σ values in units of *µ*m is described in Sec. “Conversion of arbitrary length unit to microns within the MC-TAD algorithm” in the S1 Text.

The same MC-TAD algorithm can be applied to higher resolutions (smaller bin size), see Sec. “Calculating *σ*_(3_*_×_*_3_*_×_*_3)_ and *σ*_(4_*_×_*_4_*_×_*_4)_” in the S1 Text. In the following, we present the results for *N* = 3 and *N* = 4, representing 27- and 64-bin TADs, respectively. Namely, for the case *N* = 3, which corresponds to the ∼ 4-kb chromatin resolution, we obtain *σ*_(3_*_×_*_3_*_×_*_3)_ = 0.33. For the currently highest experimentally achievable 1–2-kb chromatin resolutions [18], roughly corresponding to the case *N* = 4 (64-bin TAD), the estimated value *σ*_(4_*_×_*_4_*_×_*_4)_ is:

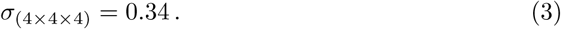

### Resolution matching between models of chromatin

In chromatin models, where the smallest resolved portion of the chromatin (bead) corresponds to TAD, the conformational heterogeneity at genomic separations smaller than TAD is exactly zero. At the resolution limit of these models the spread Σ of *R_s_*curves is practically zero (see Fig 7a, models of Tolokh et al., 2023 and Li et al., 2017). For these models, in order to estimate conformational heterogeneity at higher resolutions (e.g., within a TAD or even between TADs), we use the approximate chromatin conformations within a TAD generated by the MC-TAD algorithm. This approach allows us to approximate conformational heterogeneity that would be seen if the minimal spatial unit, a bead, were smaller than that of the TAD. This heterogeneity mimics the spread of ⟨*R_s_*⟩ curves that would effectively correspond to a higher resolution model. In the case of the two low resolution models considered that have average X chromosome TADs of 118 kb, and we can *up-convert* them to the resolution of 14 kb, using Σ_(s=118kb)(14kb)_ = *C_conv_* × *σ*_(2_*_×_*_2_*_×_*_2)_, where the parameter *C_conv_*converts the dimensionless *σ*_(2_*_×_*_2_*_×_*_2)_ to units of microns, appropriate for ⟨*R_s_*⟩, see S1 Eq 7 in the S1 Text.

The *resolution up-conversion* procedure, Fig 4, works as follows. We generate random numbers (the number of random values is equal to the ensemble size) weighted according to the distribution of spatial distances of the paths and shift up and down the original ⟨*R_s_*⟩ curve (from the model trajectory) with these numbers to emulate the spread of the curves with the same dispersion. Specifically, for up-converting 118-kb resolution models to 14-kb resolution, we generate the spread of ⟨*R_s_*⟩ curves at *s* = 118 kb approximated by a normal distribution with Σ_(s=118kb)(14kb)_ and *µ* =0. This approach assumes that chromatin within each TAD equilibrates on relatively short time scales. The results of this procedure are shown in Figs 7a and 7b, which represent the sets of ⟨*R_s_*⟩ dependencies before and after applying the up-conversion. In the plots after the up-conversion, we use a linear interpolation to connect new ⟨*R_s_*⟩ values at *s* = 118 kb to that at the new resolution size, e.g., ⟨*R_s_*⟩ = 0.1 *µ*m at *s* = 14 kb in Fig 7. In *Relative* ⟨*R_s_*⟩ plots the interpolation is to 1.00 at the resolution limit, e.g., at *s* = 2 kb in Fig 10. The purpose of the interpolation is only to guide the eye: the values of the connecting data points are not used for any further analysis.

Approximating the *Relative Conformational Heterogeneity* by a linear interpolation in Fig 9 is suggested by the *Relative Conformational Heterogeneity* dependence seen in the same figure for 10-kb resolution models of Ulianov et al., 2021, which are based on single cell Hi-C data. For up-converting the 118-kb resolution models of Tolokh et al., 2024 to 2-kb resolution, described in Results and discussion, we follow the above procedures with Σ_(s=118kb)(2kb)_ = *C*_conv_ × *σ*_(4×4×4)_.

Note that the up-conversion is most relevant only at the genomic distances equal or close to the resolution limit of the model, which is ∼100 kb in the case of Li et al., 2017 or Tolokh et al., 2023 models. Already at twice that distance, at ∼200 kb, the underlying polymer model itself produces conformational heterogeneity on its own, which grows rapidly with genomic separation. Thus, the impact of the up-conversion and its inherent approximations, such as the use of computationally manageable “two-cube” MC-TAD algorithm, is the strongest at the genomic separations close to the resolution limit of the model, and rapidly diminishes with the separation. This expectation is borne out by our results, see below.

## Results and discussion

The main goal of this work is to develop and test new metric – *Conformational Heterogeneity* (*C.H.*) – which allows one to quantify cell-to-cell heterogeneity of the interphase chromatin structures. To illustrate utility of the new metric we employ it to compare chromatin models resulting from three distinct modeling approaches that use either bulk Hi-C or single-cell Hi-C data for the fruit fly genome as the input for modeling. We also apply it o explore the effect of lamins depletion.

To mitigate possible effect on these comparisons that stem from different spatial/genomic resolutions of different chromatin models, we have developed an algorithm that allows “up-conversion” of the lower resolution chromatin structures to higher resolution ones, see Methods and Fig 7.

### Structural heterogeneity of chromatin depends on the model resolution

A simple illustration of how conformational heterogeneity can depend on the chromatin model resolution is presented in Fig 6: the two chains of beads representing chromatin conformations can look very different at higher resolution (small green spheres), while being exactly the same at a lower resolution (large white spheres). Indeed, no matter how much of conformational heterogeneity may be seen at a higher resolution, the heterogeneity can all but disappear or be much smaller at a low resolution due to coarse-graining. As a trivial example: all nuclei are the same if represented by a single spherical bead the size of the nucleus, regardless of the intricate details of the chromatin packing inside each nucleus. Therefore, to compare two or more models of chromatin packing at different resolutions, an additional step of “resolution matching” may be desirable. Arguably the most straightforward approach could be coarse-graining of the high resolution model(s) to match the level of detail of the lowest resolution model. An obvious advantage of this option is its simplicity. However, that simplicity comes at a price of information loss. Among the three models considered in this study (Methods), the model with the highest (10-kb) resolution, the Ulianov et al., 2021 model [28], is the one that is, arguably, closest to the experimental single cell (scHi-C) maps. To keep this highest resolution model intact, we propose a procedure to “up convert” lower resolution models to approximately match the 10-kb resolution of Ulianov et al., 2021 model, see Methods. As we shall see later, an additional benefit of introducing the resolution enhancement procedure is the ability to explore the limit of the highest possible scHi-C resolution on the heterogeneity of conformational ensembles of single cell nuclei.

**Fig 6.**
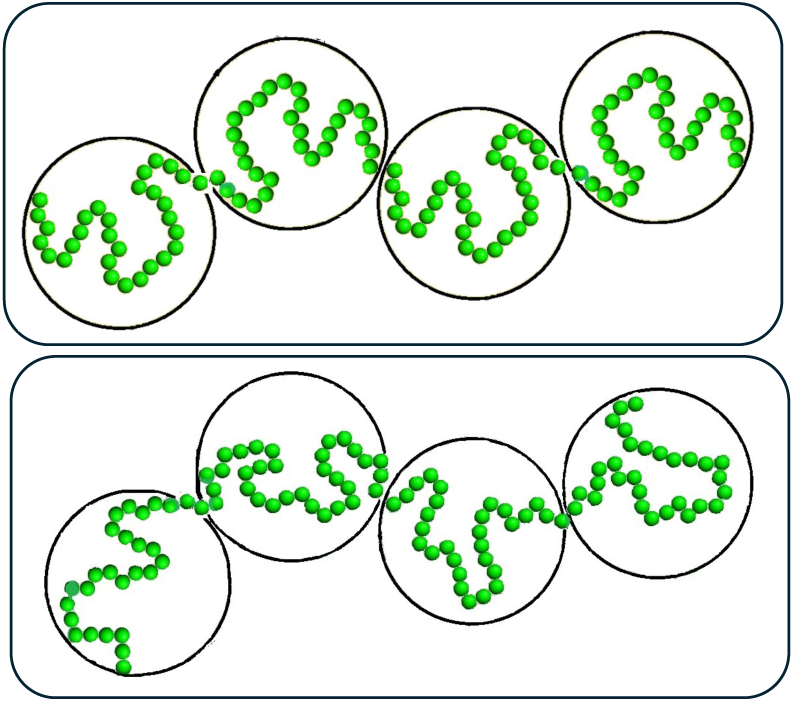
Dependence of the heterogeneity of 3D genome model conformations on the model resolution. The conformations of the two chains of green beads – chromatin models at a high resolution – are very different. After coarse-graining (going to a lower resolution), the two chromatin chains now represented by white spheres look exactly the same.

### *Conformational Heterogeneity*: A metric to characterize cell-to-cell heterogeneity of 3D Genome Organization

We propose to quantify heterogeneity of 3D chromatin structures across multiple nuclei configurations by a single metric that we call *Conformational Heterogeneity* (*C.H.*) We define *C.H.* as the standard deviation of the average spatial Euclidean distances ⟨*R*^(*k*)^⟩ between chromatin loci separated by genomic distance *s*:

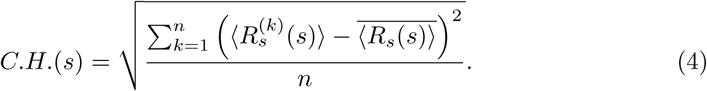

Here, each ⟨*R*^(^*^k^*^)^⟩ represents an average along *k^th^* (out of *n*) *single cell* chromatin conformation (see Methods, and S1 Eq 2 in the S1 Text), while ⟨*R_s_*⟩ refers to the average value of the ⟨*R*^(^*^k^*^)^⟩ for the entire ensemble of *n single cell* chromatin conformations (replicas). By construction, the metric aims to average out conformational differences within each given nucleus, to focus on the cell-to-cell variability. That is *C.H.*(*s*) would yield zero for a hypothetical case of identical nuclei, even if the chromatin conformations within the nucleus were highly variable.

### Visualizing variations of ⟨***R_s_***⟩ curves for the three models of Drosophila X chromosome

To begin, we have calculated the averaged Euclidean inter-loci distances ⟨*R_s_*⟩ for the set of *n* = 18 chromatin conformations of X chromosome extracted from the original Tolokh et al., 2023 modeling results, and for the sets of *n* = 20 chromosome X conformations representing both the Ulianov et al., 2021 and Li et al., 2017 chromatin models. The dependencies of ⟨*R_s_*⟩ vs. genomic inter-loci distance *s* for each of these three sets are presented in Fig 7a (the left three panels).

**Fig 7.**
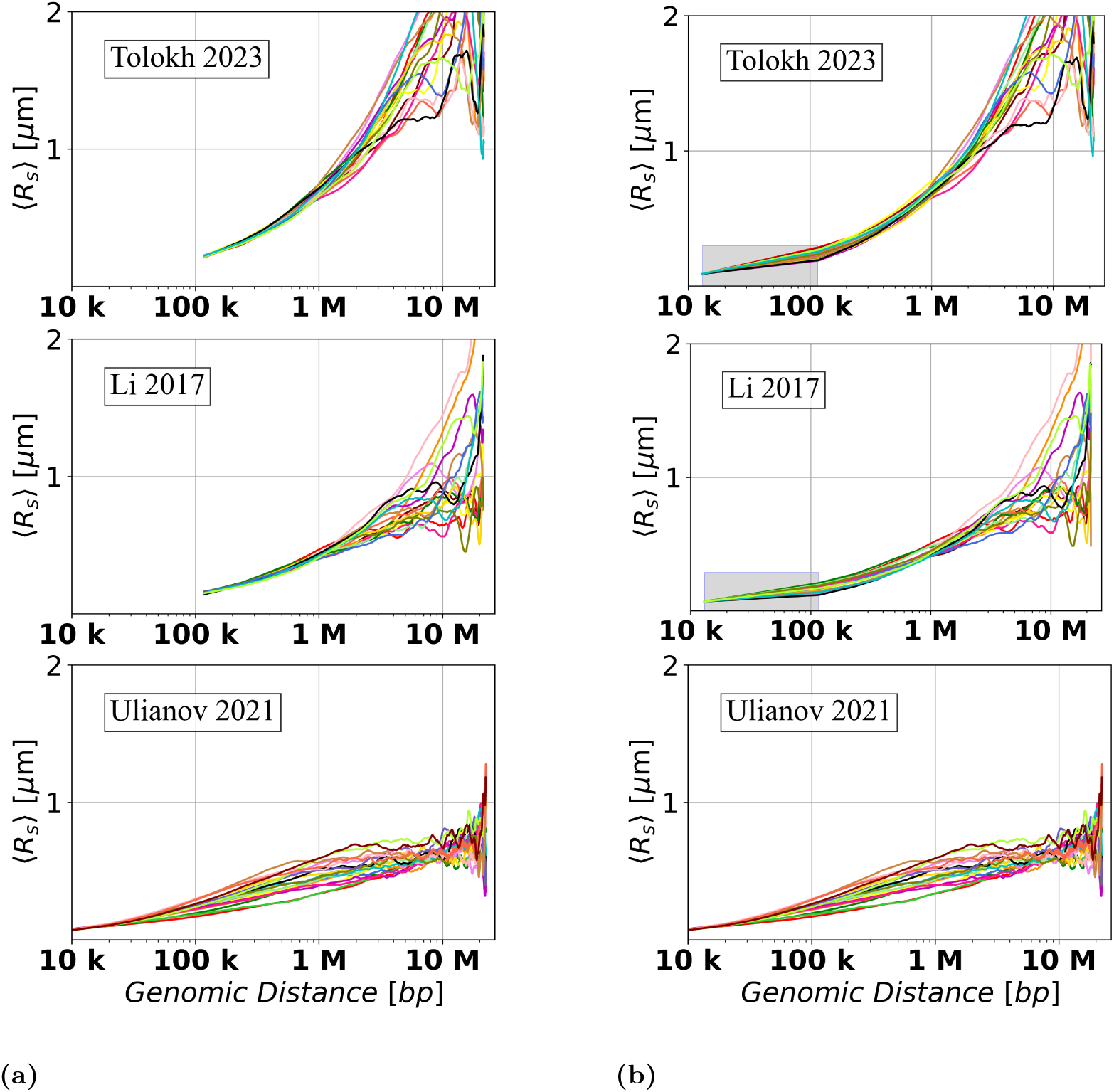
Dependencies of the averaged Euclidean inter-loci distances ⟨*R_s_*⟩ on genomic separation *s* for the three models of the X chromosome in *Drosophila* nuclei (a) before and (b) after the resolution up-conversion for the Tolokh et al., 2023 and Li et al., 2017 models to match the 10-kb resolution of Ulianov et al., 2021 models, see Methods. The Ulianov et al., 2021 models do not require up-conversion: the right panel is identical to the left panel, and is shown here for the ease of visual comparison. Each curve represents ⟨*R_s_*(*s*)⟩ dependence for one particular *single cell* X chromosome conformation. In (b), a linear interpolation is used to guide the eye in the shaded regions, see Methods.

To facilitated comparison between these three models, originally created at different chromatin resolutions, we have applied the resolution up-conversion procedure (see Methods) to the two lower resolution models. The resulting ⟨*R_s_*(*s*)⟩ “resolution-matched” dependencies are presented in Fig 7b (the three right panels).^1^

The spread of the up-converted ⟨*R_s_*⟩ dependencies in Fig 7b at around 100 kb (average genomic TAD size) is similar to the spread of these dependencies derived from Ulianov et al., 2021 10-kb resolution models without the up-conversion.

### *Relative Conformational Heterogeneity*: a detailed analysis

As is evident from the results presented in Fig 7, the absolute values and the spread of the ⟨*R_s_*⟩ curves strongly depend on the genomic distance *s* between the loci, which makes it difficult to compare the *C.H.* values within the same chromatin model for a very wide range of different *s*. A related problem arises when comparing *C.H.* values between different chromatin models. In our case, the simulations used to generate the Ulianov et al., 2021 chromatin models employed arbitrary units of distance, which are not directly comparable to micron units used by the two other models considered here. To address these issues, we define a new, normalized, and dimensionless quantity:

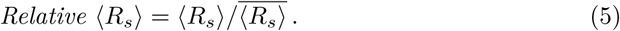

While several normalization approaches have been applied in biological experiments [46, 66], we opt for this method that adjusts each value by the overall mean ratio at each specific genomic separation. This approach is not only simpler and more interpretable than the quantile normalization used in [66], but also preserves the distribution of ⟨*R_s_*⟩ values across single cells. In contrast, quantile normalization assumes that all ⟨*R_s_*⟩ values follow the same distribution, which may not be realistic as seen from Fig 8.

**Fig 8.**
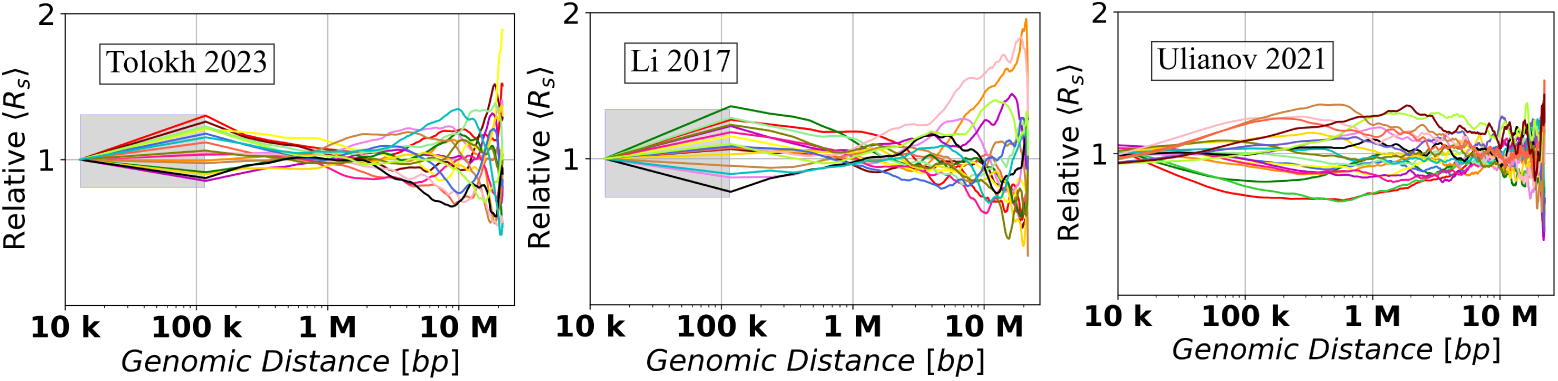
Dependencies of the *relative* Euclidean inter-particle distances (*Relative* ⟨*R_s_*⟩), calculated using Eq 5, on the genomic separation *s* between the polymer beads for each of the three chromatin models on a log-scaled x-axis. In the shaded regions a linear interpolation is used to guide the eye, see Methods.

Fig 8 shows the dependencies of *Relative* ⟨*R_s_*⟩ values on the genomic distance *s* calculated from the up-converted ⟨*R_s_*⟩ curves presented in Fig 7b for the three different models of X chromosome in *Drosophila*. To characterize the spread of *Relative* ⟨*R_s_*⟩ curves, we introduce the corresponding quantity – *Relative Conformational Heterogeneity* (*Relative C.H.*):

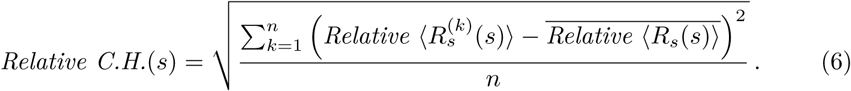

Note that just like the absolute C.H. of Eq 4, the *Relative C.H.* focuses on conformational variability *between* nuclei in an ensemble. The *Relative* ⟨*R_s_*⟩ curves, Fig 8, allow one to estimate the dependencies of the *Relative C.H.* values, Eq 6, on the genomic separation *s*. The corresponding *Relative C.H.* dependencies are shown in Fig 9a, while the bar chart in Fig 9b shows the *Relative C.H.* values at the three specific inter-loci genomic separations *s*.

**Fig 9.**
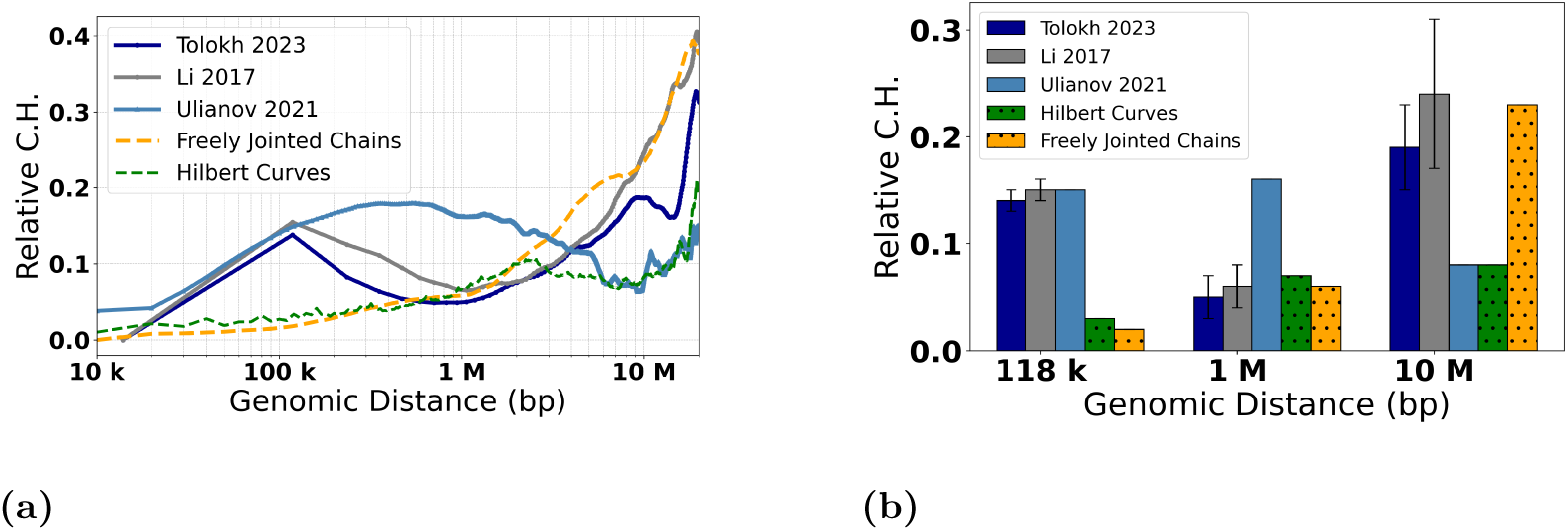
*Relative C.H.* values, Eq 6, for the three models of 3D chromatin organization in X chromosome of *Drosophila* at approximately 10-kb resolution, as well as two highly simplified polymer chain models based on the Hilbert Curve (space-filling fractal) and the Freely Jointed Chain (FJC), of the same genomic length as the X chromosome, see Methods. **(a)** The dependencies of the *Relative C.H.* on the genomic separations *s*. For Li et al., 2017, and Tolokh et al., 2023 models, a linear interpolation is used from 14 kb to 118 kb; for the FJC *Relative C.H.* ≈ 0.57 in the limit of genomic separation *s* → ∞, see Methods. **(b)** Comparison of the *Relative C.H.* values for three specific genomic separations for each of the three chromatin models. The error bars represent the standard deviations estimated using 10 sets of chromatin conformations for the Li et al., 2017 and Tolokh et al., 2023 models.

To put these model-specific *Relative C.H.* numbers in perspective, we compare them with the corresponding *Relative C.H.* of two highly simplified “generic”, but distinctly different, models of chromatin packaging [42]: ensembles of space-filling fractal curves (approximate orientation-random Hilbert curves), and that of the freely jointed chains (FJC), see Methods. As expected, all the *Relative C.H.* curves converge to essentially zero^2^ at the lower end of the genomic distance range – the resolution limit of the models. As expected, the *Relative C.H.* values for all the models increase as one moves away from their resolution limit to 118 kb, which is the average TAD size for the three models of X chromosome. Note that ∼10 kb is the resolution limit for all three models of X chromosome (with the up-conversion for Tolokh et al., 2023 and Li et al., 2017 models). At the opposite end of the range, near *s* = 20 Mb, all of the *Relative C.H.* curves exhibit a sharp rise, which can be explained as follows. By the definition of ⟨*R_s_*⟩ (see S1 Eq 3 in the S1 Text), it is an average over all of the bead-bead pairs separated by the same genomic distance *s* within each cell: there are many such pairs for the mid-range distances, which, as intended, dampens the effect of random variations within each nucleus. But only a few pairs exist at the upper end of the range, e.g., just one for the maximum *s*. In this regime, the influence of statistical fluctuations becomes highly noticeable. Finally, we notice that, compared to the simplified polymer models, for all three realistic models of X chromosome in Fig 9, the *Relative C.H.* is considerably higher, up to ∼500 kb genomic separation. What it likely means is that the cell-to-cell variability of real chromatin structures at relatively small and medium scales is higher than what can be inferred from highly simplified polymer models such as FJC or an idealized space-filling fractal.

Next, we focus on the three models of *Drosophila* X chromosome, and the details of the corresponding *Relative C.H.* dependencies presented in Fig 9, noting that they are derived from the X chromosome models generated using three completely different computational approaches. At ∼100 kb of genomic separations, all three models have approximately the same *Relative C.H.*, suggesting that the resolution “up-conversion” procedure applied to the Tolokh et al., 2023 and Li et al., 2017 models produces chromatin conformations that are as structurally heterogeneous from cell to cell as those derived from the scHi-C maps with 10-kb resolution.

As the genomic separation increases further, the behavior of *Relative C.H.* of the chromatin models derived from the bulk Hi-C map (Li et al., 2017 and Tolokh et al., 2023 models) begins to differ markedly from those based on the scHi-C maps (the Ulianov et al., 2021 models). From 118 kb to about 1 Mb, the first two models demonstrate a steady decrease of the *Relative C.H.* values, reaching a minimum, 0.05, at about 1 Mb, while Ulianov et al., 2021 models demonstrates an opposite behavior, reaching the maximum of their *Relative C.H.* value of approximately 0.18 at about 0.8 Mb.

The opposite behavior is also observed in the genomic separation range from 1 Mb to 10 Mb: the two bulk Hi-C based models exhibit a steady increase of their *Relative C.H.* values to about 0.2, while the models derived from the scHi-Cs (the Ulianov et al., 2021) show a decrease of the *Relative C.H.* to a minimum, of 0.05, at 10 Mb. At larger genomic separations, the bulk Hi-C based models continue to demonstrate larger *Relative C.H.* values than the scHi-C based models.

We suggest that the described above major differences in the *Relative C.H.* behavior between the bulk vs. single-cell Hi-C based chromatin models are related to several factors. One of them, most relevant at 0.2 - 2 Mb separations (2 to 20 TADs), is the presence of relatively weak (interaction well depth 0.5 to 1.5 kT) but frequent TAD-TAD interactions at these genomic separations in the Tolokh et al., 2023 dynamic chromatin models. These interactions were optimized to reproduce the experimental bulk Hi-C map, which exhibits numerous TAD-TAD contacts at these genomic separations. The same experimental bulk Hi-C map is used to select chromatin conformations generated in the Li et al., 2017 modeling approach, resulting in approximately the same *Relative C.H.* (see Fig 9). However, most of these TAD-TAD contacts are absent in a small number (20) of the scHi-C maps used in the Ulianov et al., 2021 modeling approach. We speculate that using only a limited number of the inter-TAD restrains corresponding to a very limited number TAD-TAD contacts in each of these 20 scHi-C maps results in a larger spatial heterogeneity of the conformations at these (0.2 - 2 Mb) genomic separations in the 3D models of X chromosome in the modeling approach of Ulianov et al., 2021. This increased *Conformational Heterogeneity* leads to a wider spread of the *Relative* ⟨*R_s_*⟩ curves compared to the two other models (Fig 8), leading to larger *Relative C.H.* values (Fig 9) at these genomic separations.

Regarding the opposite behavior of the Tolokh et al., 2023 and Li et al., 2017 models vs. Ulianov et al., 2021 models at 10 Mb and larger genomic separation, Fig 9, we suggest the following explanation: the difference could be related to the restrictions on the simulation volume in these different models. Note that spatial confinement is a major determinant of many aspects of the chromosome organization in 3D space [13, 67, 68]. Namely, the conformational models of Li et al., 2017 and Tolokh et al., 2023 restrict the size of X chromosome to roughly 4 micrometers – the size of the fruit fly nucleus, while in the Ulianov et al., 2021 models the whole simulated system is retained within a cubic cell of only about 2 micrometers (22 DPD a.u.). Naturally, the resulting 3D structures of X chromosome are relatively more compact (see S1 Fig G and S1 Table F in the S1 Text) and, therefore, are expected to exhibit more limited heterogeneity at these (larger than 10 Mb) genomic separations.

To verify that the lower *Relative C.H.* in Ulianov et al., 2021 model at 10 Mb (lower compared to the models of Tolokh et al., 2023 and Li et al., 2017) is not an artifact of the resolution up-conversion in these two originally lower resolution models, we checked the *Relative C.H.* values at 10 Mb in these Tolokh’s and Li’s models without the resolution up-conversion. The resulting values are 0.18 for Tolokh et al., 2023 and 0.24 for Li et al., 2017 models, which are virtually the same as those after the up-conversion: 0.19 and 0.24, respectively, Fig 9b (and S1 Table D in the S1 Text). Thus, with or without the up-conversion, the *Relative C.H.* in Tolokh et al., 2023 or Li et al., 2017 models at 10 Mb remains two to three times higher than that one in Ulianov et al., 2021 models.

In summary, the results presented in Fig 9 demonstrate a very similar *Conformational Heterogeneity* for Tolokh et al., 2023 and for Li et al., 2017 models of fruit fly X chromosome, which are based on bulk (ensemble averaged) Hi-C maps; the heterogeneity in Ulianov et al., 2021 models, based on a relatively small set scHi-C maps, are significantly different.

### *Relative C.H.* at the resolution extrapolated to 2 kb

Here, we ask what might be the *Relative C.H.* of 3D genome organization in *Drosophila* nucleus at approximately 1–2 kb resolution – arguably the highest resolution possible for present day Hi-C experiments [18]. The extrapolation is exemplified for the models from Tolokh et al., 2023, and is based on Σ_(s=118kb)(2kb)_ = 0.05 *µ*m, see Methods, and further details in “Extension to higher resolution” in the S1 Text. The results are summarized in Fig 10 and Table 1.

**Fig 10.**
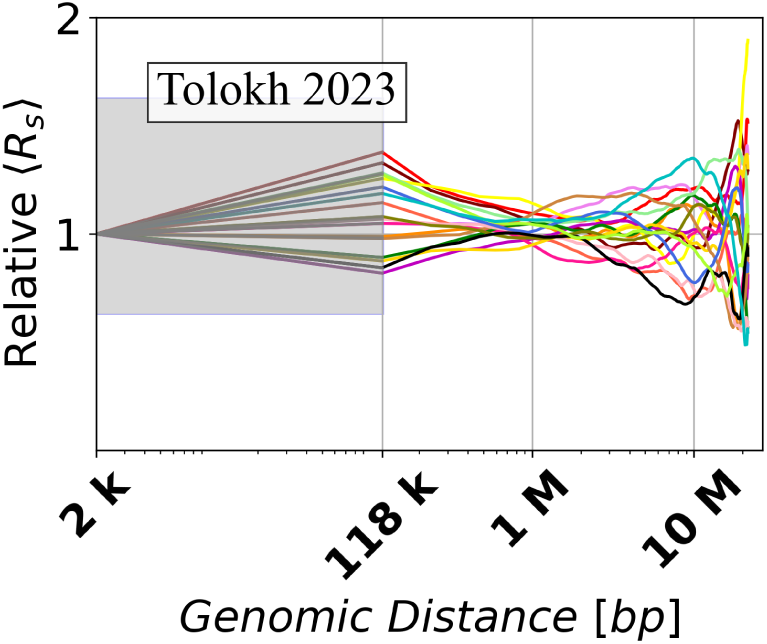
Extrapolation of the ensemble X chromosome conformations from Tolokh et al., 2023 models to 2-kb resolution, corresponding to Σ_(s=118kb)(2kb)_ = 0.05 *µ*m. Shown are the dependencies of the *relative* mean Euclidean inter-particle distances on the genomic separation *s* between the polymer beads calculated using Eq 5. In the shaded region a linear interpolation is used to guide the eye, see Methods.

Comparing these *Relative C.H.* values with the corresponding ones for the same models at 14-kb resolution, see Results and discussion and S1 Table D in the S1 Text, one can see only a very slight increase, from 0.14 to 0.17, of the *Relative C.H.* at 118 kb, with no discernible change at 1 Mb and 10 Mb. This small difference means that 14-kb resolution is good enough for describing conformation heterogeneity at much larger scales of genomic inter-particle separations within the types of chromatin models considered here. For completeness sake, an extrapolation to the theoretical limit of single base-pair resolution is presented in the S1 Text, where one can see a significant, ∼3 fold increase of the *Relative C.H.* at 118 kb, a moderate (∼2 fold) increase at 1 Mb, but still no change at 10 Mb.

### The effect of lamins depletion on the structural heterogeneity

In eukaryotic cells the internal surface of the nuclear envelope is lined by the nuclear lamina – a network of lamins and associated proteins [69, 70]. Lamins are an important determinant of nuclear architecture and gene expression [70]; mutations that disrupt normal functions of lamins play a central role in a diverse set of genetic disorders known as laminopathies, which have symptoms that range from muscular dystrophy to neuropathy to premature aging [71]. Lamina-associated domains (LADs) are regions of the genome that frequently interact with the nuclear lamina, the layer just inside the nuclear envelope. These interactions help anchor portions of the genome near the nuclear periphery, and are correlated [49] with transcriptionally inactive chromatin. Although *Drosophila* LADs are dynamic and can detach and reattach during interphase [13], their affinity for the nuclear lamina plays a stabilizing role in how chromatin is organized inside the nucleus [72], including stabilization of the chromosome territories [37].

Depletion of lamins, caused either by a lamin mutation [73] or knockdown [13, 74], leads to re-arrangement of chromatin, but, surprisingly, the effect is barely noticeable from the corresponding Hi-C maps [13, 74]. Here we ask how lamins depletion, which abolishes the affinity between LADs and the nuclear envelope (see Methods), affects the *Relative C.H.* of the X chromosome in fruit fly nuclei. A clear trend seen in Fig 11 is that for nearly all genomic separations, the structural cell-to-cell heterogeneity of the lamins depelted nuclei is greater than that of the wild type. The trend is consistent with the accelerated loss of chromosome territories in lamins depleted nuclei [37]. An important methodological implication is that *Relative C.H.* is sensitive enough to characterize this nuanced difference. A potential biological significance of this finding is that cell functions that depend on chromatin structure, such as transcription levels, might be more variable in lamins depleted nuclei.

**Fig 11.**
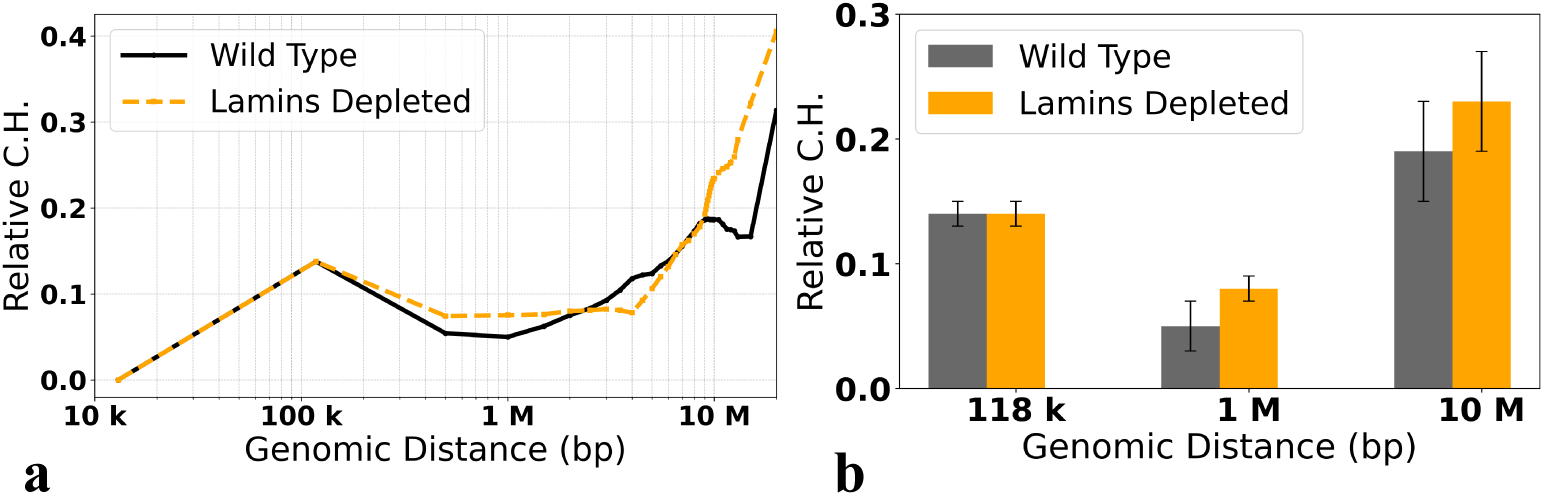
*Relative C.H.* values, Eq 6, for the Wild Type and Lamins depleted models of 3D chromatin organization in X chromosome of *Drosophila* (Tolokh et al., 2023). **(a)** The dependencies of the *Relative C.H.* on the genomic separations *s*. **(b)** Comparison of the *Relative C.H.* values for the three specific genomic separations. The error bars represent the standard deviations estimated using 10 sets of chromatin conformations, each for the Wild Type and Lamins depleted nuclei.

## Conclusion

In this work we have explored, computationally, how cell-to-cell *Conformational Heterogeneity* of genome organization in 3D space varies across three different models of fruit fly nucleus, focusing on X-chromosome for concreteness. We have also explored the effect of nuclear lamins depletion on the *Conformational Heterogeneity*. The *Conformational Heterogeneity* naturally arises from differences in chromatin folding patterns from cell to cell of the same tissue, as well as from the time evolution of chromatin conformations. Thus, to characterize the cell-to-cell variability of chromatin in 3D, one needs to characterize the distribution of the structural states of the chromatin chain in space, focusing on how it varies between the nuclei. Our first result is a general metric that can be used to quantify *Conformational Heterogeneity*: we have defined it for an ensemble of single cells, as the standard deviation of the distribution of the averaged (over each cell) Euclidean inter-loci distances, ⟨*R_s_*⟩, calculated for a set of conformations/states of the chromatin chain for a given genomic separation *s* between two loci. As defined, *Conformational Heterogeneity* is simply the second moment of the corresponding distribution – a single number for a given genomic scale, *s*, with a clear interpretation. By construction, the metric aims to average out conformational differences within each given nucleus, to focus on the cell-to-cell variability. That is it would yield zero for a hypothetical case of an ensemble of identical nuclei, even if the chromatin conformations within the nucleus were highly variable. We also introduced *Relative Conformational Heterogeneity* – a dimensionless version of the metric, derived from the distribution of ⟨*R_s_*⟩ normalized by their ensemble-averaged value. We have argued that the relative metric is better suited for comparing different models of chromatin across a range of genomic distances, as well as across different models of chromatin.

We have explored *Relative Conformational Heterogeneity* for three distinct, existing models of 3D chromosome organization in fruit fly nuclei – the goal was to identify common general trends while also exploring potential differences. The first two models, Li et al., 2017 [41] and Tolokh et al., 2023 [13], aimed to approximate the bulk Hi-C map from the experiment, but in very different ways. Li et al. [41] aimed to obtain a virtually exact match to the HiC map for a large, time-independent ensemble of selected conformations of chromosomes, while Tolokh et al. [13] used a polymer physics model to describe time-evolution of chromosomes in the population of nuclei, with the match to the same experimental bulk Hi-C map being more approximate than in Li et al. [41] modeling approach. In contrast, the third set of chromosome models (Ulianov et al., 2021 [28]) was derived from a limited number of single-cell Hi-C maps [28], which made it a unique point of reference to compare to. Note that even hypothetical models constructed as perfect fits to perfect, but different types of Hi-C reference data, may yield different structural ensembles. Furthermore, the same bulk Hi-C map can originate from substantially different populations of single-cell Hi-C maps, see the S1 Text for a highly simplified example. To better understand the behavior of these realistic models of chromatin, we have also compared the corresponding *Conformational Heterogeneity* to those of two highly simplified polymer models – ensembles of space filling fractals and freely jointed chains.

Application of the *Relative Conformational Heterogeneity* metric to each of the above models has led to several findings.

First, we have found that *Conformational Heterogeneity* of chromatin depends strongly on the model resolution, with higher resolution models being more conformationally heterogeneous in general. In retrospect, this finding should not come as a surprise: a highly coarse-grained model of the same polymer is not expected to show greater variety of structures compared to its finer representation: in fact, any model is expected to show near zero *C.H.* right at its resolution limit. We have seen just that with all the models; all of the models, including the two two highly simplified polymer models, show very similar trend of increasing structural heterogeneity starting from zero or near zero at the model resolution limit (10 kb in our case) to about 100 kb in genomic separation, which is the average TAD size for the realistic models of X chromosome used here. Another general trend seen for all the models is the sharp increase of *C.H.* at the largest genomic separations, which probes cell-to-cell variability of Euclidean distance between pairs of loci near the opposite ends of the chromosome.

Second, our work has made it possible to explore the relationship between *Conformational Heterogeneity* and model resolution quantitatively, within a certain set of assumptions. Although increasing model resolution has a dramatic impact on the *Conformational Heterogeneity* near the genomic separation *s* corresponding to the resolution limit of the model, the impact decreases for larger genomic separations. For example, extrapolating model resolution from approximately 10 kb to the current experimental Hi-C data limit of 1 kb has little effect on the *Conformational Heterogeneity* beyond *s* ≈ 100 kb. Reaching a hypothetical single base-pair resolution would have a much larger effect, but it still disappears at the largest genomic separations of 10 Mb.

Third, for the two realistic models that aim to approximate the bulk Hi-C map of fruit fly, the behavior of *Conformational Heterogeneity* is nearly the same across all genomic distances from 10 kb to 10 Mb, apparently insensitive to different specifics of each model, as long as they both approximate the reference bulk Hi-C map well (in the case of Li et al., 2017 model the match to the Hi-C map is essentially exact by construction). At the same time, the realistic model (Ulianov et al., 2021) aimed to reproduce available single-cell Hi-C data showed significant, non-monotonic differences (in the behavior of the *Conformational Heterogeneity* metric) from the first two models beyond 118 kb (TAD size) genomic separation, immediately suggesting that our new metric can be quite sensitive to specifics of the reference Hi-C data. We have proposed a plausible explanation for the observed differences from a polymer modeling perspective. Based on this analysis, we propose to explore the possibility of inclusion of bulk Hi-C data into model construction procedure for models of 3D chromatin that utilize necessarily limited single-cell Hi-C data. We argue that a reasonable agreement with the bulk Hi-C map might increase the likelihood that the single-cell Hi-C based approach generates chromatin conformations that reproduce not only the limited set of loci contacts from available single-cell Hi-C data, but also the much larger set of different inter-loci contacts, with proper statistics. We speculate that incorporating an additional fit to the appropriate bulk Hi-C map, *e.g.*, in the form of additional inter-TAD restraints on chromosome conformations, might improve realism of such models: these additional restrains can be based on the TAD-TAD contacts missing from the available single-cell Hi-C data. In the future, as the single-cell Hi-C technique matures to produce much larger ensembles that represent real populations more faithfully, these additional restrains can be dropped. In principle, a modeling approach that is able to reproduce a large, statistically significant ensemble of single-cell Hi-C maps, which faithfully represents the true distribution of chromatin states in a cell population, should be superior to a modeling approach that only reproduces a single bulk Hi-C map. From this perspective, the pioneering single-cell Hi-C based modeling approach of Ulianov *et al.* [28] represents an important first step in the highly promising direction.

While the focus of this work is almost exclusively methodological, we have shown how the new metric of cell-to-cell conformation heterogeneity can be used to gain insights into a biologically relevant problem of the effect of nuclear lamins depletion of conformational heterogeneity of the X chromosome in fruit fly. We find that for nearly all genomic separations, the structural heterogeneity of the lamins depelted nuclei is greater than that of the wild type; the trend is consistent with the accelerated loss of chromosome territories in lamins depleted nuclei [37]. Based on this finding, we have made a prediction that cell functions that depend on chromatin structure, such as transcription levels, might be more variable – “noisy” – in lamins depleted nuclei. While these specific predictions are made based on the models of fruit fly nuclei, we speculate that increased cell-to-cell structural heterogeneity of chromatin conformations, and the associated biological consequences, might extend beyond fruit fly: this is because as in fruit fly, lamins depletion in mammalian nuclei also promotes deterioration of chromosome territories [75].

Beyond the initial exploration of the new metric for *Conformational Heterogeneity* of 3D chromatin conformations presented here in the context of three specific models, several future directions are worth mentioning. It will be interesting to explore how *Conformational Heterogeneity* evolves in time in the interphase, and, more generally how *Conformational Heterogeneity* may evolve as the cell goes through rounds of cell divisions, that is, as the cell ages. The metric can also be employed to investigate how *Conformational Heterogeneity* changes in diseased vs. normal cells, e.g., in cancer or other disorders linked to notable chromatin re-arrangements [76–78]. It is likely that the *Relative Conformational Heterogeneity* – made dimensionless by construction – is suitable for comparisons across tissue types and, possibly, even among different organisms. We stress that the current version of the metric is developed for a single contiguous piece of the genome, e.g., a locus or a single chromosome – a limitation that might be overcome in the future. Also, as defined, *Conformational Heterogeneity* represents only one moment of the corresponding complex distribution of structural states. In the future, it would be interesting to explore the possibility to characterize cell populations by higher moments of ⟨*R_s_*⟩ distribution. Since many computational models of the 3D chromatin organization automatically provide a large set of “single cell” structural snapshots, we expect that the methodology developed in this work can be easily adopted for these other studies, beyond fruit fly.

## Data Availability

The snapshots of the 3D conformations of X chromosome along with collection of codes and scripts needed to compute *Relative* ⟨*R_s_*⟩, *C.H.* and *Relative C.H.* from this data, as well a the script used to generate the approximate Hilbert curves, are available on GitHub: https://github.com/Onufriev-Lab/hi-c_model_validation. The data corresponding to the Monte Carlo simulations is available at the same link, with full documentation available in the Supplementary folder. We have followed the general guidelines outlined in *“Ten simple rules on how to create open access and reproducible molecular simulations of biological systems”* [79].

## Supporting information

**S1 Text. Supplementary Methods.**

## Acknowledgment

We would like to thank Alexandra Galitsyna and Pavel Kos for many helpful discussions and Frank Alber for providing a whole set of genome structures referenced in [41]. SM would like to express her heartfelt gratitude to the late Professor [Jie Liang], whose mentorship and dedication were instrumental in shaping her interest in the field of 3D genome. The authors acknowledge Advanced Research Computing at Virginia Tech for providing computational resources and technical support that have contributed to the results reported within this paper (URL: http://www.arc.vt.edu). Partial support from the National Institutes of Health, R01 GM144596 to A.V.O., is acknowledged.

## Footnotes

1 In the case of Li et al., 2017 and Tolokh et al., 2023 chromatin models, the spread of the ⟨*R_s_*⟩ curves at *s* = 118 kb, which is the genomic separation corresponding to the average genomic TAD size in X chromosome, are, strictly speaking, not zero, but very small: Σ_(_*_s_*_=118_*_kb_*_)_ = 0.0048 *µ*m and Σ_(_*_s_*_=118_*_kb_*_)_ = 0.0039 *µ*m, respectively. These spreads at the model resolution limit are not exactly zero due to fluctuation of the center-to-center Euclidean distances between the beads representing TADs. In what follows we do not discuss this minor effect, which is much smaller than the relevant cell-to-cell heterogeneity we seek to explore and characterize. Since the spread of the original (before up-conversion) ⟨*R_s_*(*s*)⟩ dependencies for the lower resolution models at their resolution limit *s* = 118 kb is very small, we assume that after up-conversion the spread at *s* = 14 kb for these two models is as small or smaller and set it to Σ_(_*_s_*_=14_*_kb_*_)_ = 0.00.

2 The spread for the space-filling fractal curves (the random-orientation Hilbert curves) is not strictly zero at the resolution limit because of the approximations used: the curve contains some diagonal segments which have larger length than point-to-point identical segments of unit length, see Fig 3. The spread of the curves for chromatin configurations of Ulianov et al., 2021 models at their resolution limit is related to the thermal motion of the beads, which produces different distributions of the bead-to-bead distances in different configurations.

