## Supplemental Info for "Quantifying Conformational Heterogeneity of 3D Genome Organization in Fruit Fly"

#### 1 End-to-End distance of polymer conformation

The end-to-end distance $R(L)$ of a polymer representing a chromatin fiber, and consisting of a chain of $L$ beads is the Euclidean distance between the centers of the initial and final beads for a given polymer conformation. For the $k$-th conformation, this distance can be expressed as:

$$R^{(k)}(L) = \left\| x_1^{(k)} - x_L^{(k)} \right\|, \qquad (1)$$

where $x_i$ are coordinates of the $i$-th bead. The genomic size of each of $L$ beads corresponds to a polymer (chromatin) resolution size.

Accordingly, the Euclidean distance between the centers of bead $i$ and bead $j = i + s$ for the $k$-th polymer conformation is:

$$R_i^{(k)}(s) = \left\| x_i^{(k)} - x_{i+s}^{(k)} \right\|. \tag{2}$$

The maximum value of $i$ for a given value of $s$ is $L - s$.

The average Euclidean distances for all possible pairs of $i$ and $j$ separated by $s$ for the $k$-th conformation of the polymer with $L$ beads is:

$$\langle R_s^{(k)}(s) \rangle = \frac{\sum_{i=1}^{L-s} \left\| x_i^{(k)} - x_{i+s}^{(k)} \right\|}{L - s}, \tag{3}$$

where the denominator $L - s$ is the total number of all possible subchains of $s + 1$ beads ($s$ can be considered as a center-to-center length of the polymer subchain).

#### 2 Generating permissible conformations within a single cube

The given section describes a method to simulate chromatin folding using a random walk model within one Topologically Associating Domain (TAD). The chromatin configurations (paths) must satisfy certain physical conditions, such as starting and ending on the faces of the cube and not crossing over a segment. To account for the possibility of long-range interactions or loops within a TAD boundary, we allow the paths to cross over a point. Hence, due to the possibility of unfilled bins, accurately determining the total count of the sample space becomes challenging. The physical conditions that permissible paths in the $N \times N \times N$ cube must satisfy are the following: (a) In the $N > 2$ case, the start and end of the path must be at the faces of the cube, rather than in the middle bins, because the path needs to continue to the second cube. (b) In the $N > 2$ case, the endpoint of the path cannot be at the starting face, unless the endpoint bin is at the edge. (c) The paths cannot cross over a segment of non-zero length. (d) The length of the path $s$ (the number of segments along the path in the $N \times N \times N$ cube) must be equal to $N^3 - 1$. (e) Due to symmetry, we only consider conformations that start from a specific face. For example, for the first of the two adjacent cubes we consider only the paths with the starting point located at the $X^-$ face (left face) of the cube, and because of directionality of the chromatin chain in X chromosome, the direction $X$- is excluded. The paths in the second cube can start from any of the second cube faces which can be in contact with the first cube, excluding its (1-st cube) $X^-$ face. These are all the faces except for the $X^+$ face of the second cube.

The above conditions and conditions described in “Methods" of the main text serve to establish a clear set of rules that a permissible path must follow. By outlining these conditions, we can ensure that the analyzed paths are valid and conform to the physical constraints of the system being studied. We remove repetitive paths generated by Monte Carlo procedure and maintain only the unique ones. After obtaining all unique permissible paths (conformations) in one cube, we call each one a *selected-path* or a *unique path*. A conformation of the chromatin with the average TAD length of about 118 kb can start anywhere on X chromosome, without losing generality.

To achieve this, a chain of beads/bins (a conformation) can start anywhere in one cube (TAD) and continues in the second, adjacent cube (TAD). This requires us to generate paths in two adjacent cubes.

### 3 Generating permissible conformations within two adjacent cubes

After generating permissible conformations in the first cube, the conformations in the second adjacent cube that continue the paths in the first cube are created in “two steps" as follows:

First, for each unique path in the first cube, which always start from the $X^-$ face of the cube, we determine its endpoint $x$-coordinate. *If this endpoint belongs to the* $X^+$ *face of the first cubes*, then, to determine permissible paths which continue in the second cube, we select the paths from the set of already generated unique paths for the first cube such that their starting point (if shifted in $X+$ direction) matches the endpoint of the path in the first cube (we call this *a comparison-path*). For each endpoint, all possible comparison paths are determined. Then, the paths for the second cube are generated by translating each comparison path by $N$ units in $X+$ direction.

Second, *if the endpoint y-coordinate of the unique path in the first cube belongs to* $Y^+$ *face*, we first generate a new set of all permissible unique paths (see conditions in Sec. 2) that start from the $Y^-$ face of the cube. Then, for each of the endpoint in the $Y^+$ face in the first cube, we determine all possible comparison paths from the new generated set of permissible unique paths. These comparison paths are then translated by $N$ units in $Y+$ direction, creating the paths in the second cube.

These two steps together allow one to generate continuous chromatin conformations in two adjacent TADs (cubes). Due to symmetry considerations, without loss of generality, the other possibilities (the location of endpoint in the first cube in its $Y^-$, $Z^+$ and $Z^-$ faces) are congruent with the original second step.

S1 Figure A shows four examples of continuous chromatin chain conformations (paths) along the two adjacent cubes with four different starting points, generated by the algorithm in each run which reached a valid path.

**Algorithm 1** Selection and Translation

**Input:** The unique paths in the first cube.

**Output:** The continuous paths (conformations) in the two cubes ($X$ and $Y$ directions).

**for** *selected_path* in *all_path* **do**

  **for** *comparison_path* in *all_path* **do**

    **if** (SPE_x = n) **then**: ▷ SPE is the end coordinate of the selected path

      Selection $(n, y, z) = (1, y, z)$

      Translation ((CPS_x) = 1, SPE_y = CPS_y, SPE_z = CPS_z ) ▷ CPS is the start coordinate of the comparison path.

    **end if**

  **end for**

  **for** *comparison_path_Y* in *all_path* **do** ▷ comparison_path_Y refer to all successful paths starting from the lower face (y=0) of the first cube and ending at any other faces.

    **if** (SPE_y = n) **then**:

      Selection $(x, n, z) = (x, 1, z)$

      Translation (SPE_x = CPS_x, CPS_y = 1, SPE_z = CPS_z )

    **end if**

  **end for**

**end for**

**S1 Figure A.** Four examples of permissible paths.

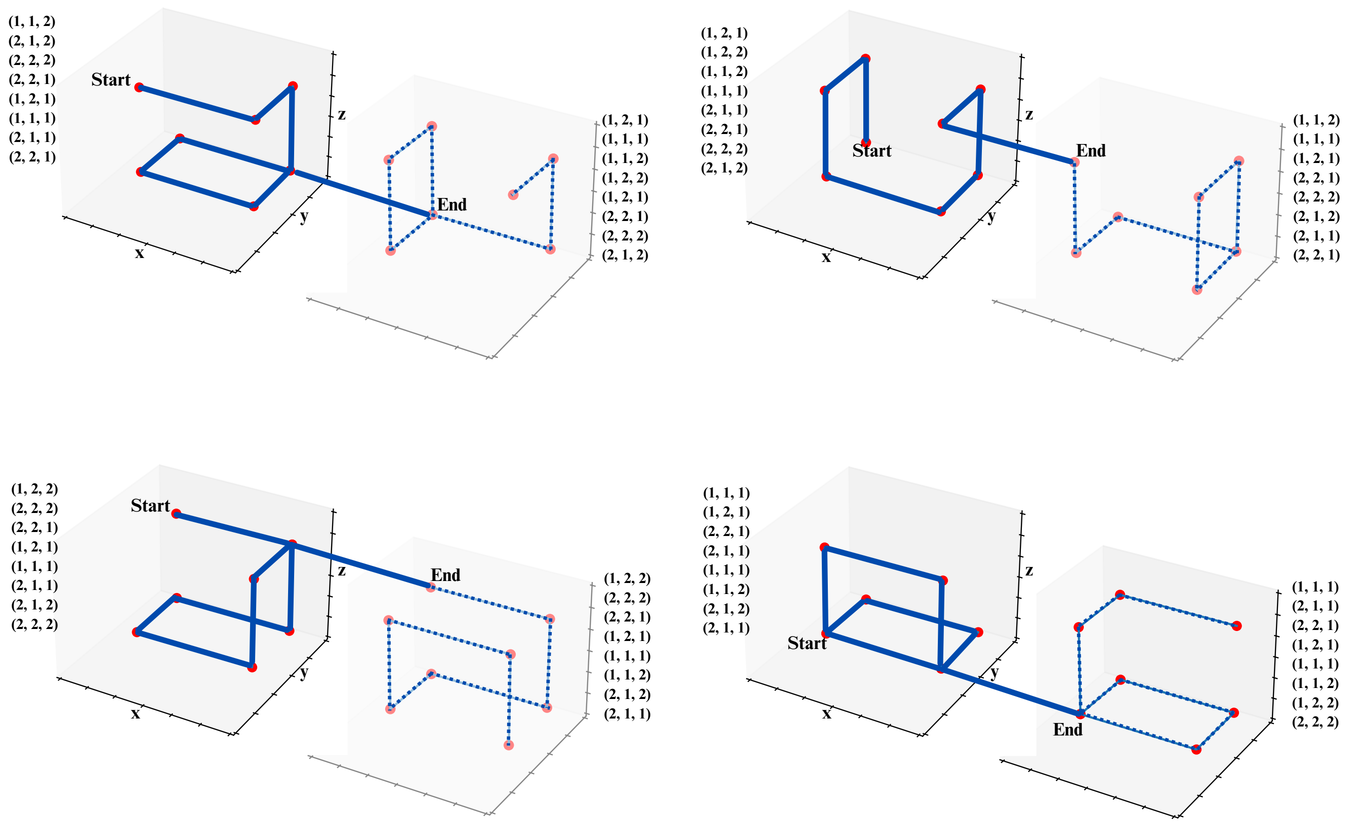


**S1 Figure A.** Four examples of permissible paths along the two adjacent cubes with four different starting points. The starting face is fixed (the left ($X^-$) face of the first cube). The cube resolution corresponds to $N = 2$ (8 bins). For the implementation convenience, we consider the size of the cube as 1.00. In addition, we use integers to label the centers of each bin. This results in the following mapping onto the (x,y,z) coordinates for the centers of bins within the unit size cube: (1,1,1) $\mapsto$ (0.25,0.25,0.25) and (2,2,2) $\mapsto$ (0.75,0.75,0.75).

##### 3.1 End-to-end distances for subchains of chromatin conformations in two adjacent cubes

We thoroughly examine all possible end-to-end distances (S1 Eq 2) for the subchains of length $s$ (number of segments) along the two connected paths in two adjacent cubes when the starting bin of a subchain is in the first cube and ending bin is in the second cube. For example, S1 Figure B shows two different subchains (separated by $s = 8$ segments) along the two different chromatin conformations in the two adjacent cubes (TADs).

The distribution of the averaged end-to-end distances (S1 Eq 3) between the chromatin loci separated by genomic distance which corresponds to the average TAD size (∼118 kb) should be within the lower and upper bounds for the possible non-averaged end-to-end distances for this genomic separation. These two limiting cases are presented in S1 Figure B for the 8-bin cubes and $s = 8$. The left panel shows the subchain conformation with the largest possible distance (for the permissible two-cube chromatin configurations at this (8 bins) resolution) – 1.58 arbitrary units. The right panel shows the subchain conformation with the smallest inter-loci distance – 0.71 arbitrary units. Since one arbitrary unit (the cube size) corresponds to the average TAD size in a chromatin model, presented in the "Methods" section of the paper, they can be easily converted to micrometers.

**S1 Figure B.** Upper and lower bounds for end-to-end distances.

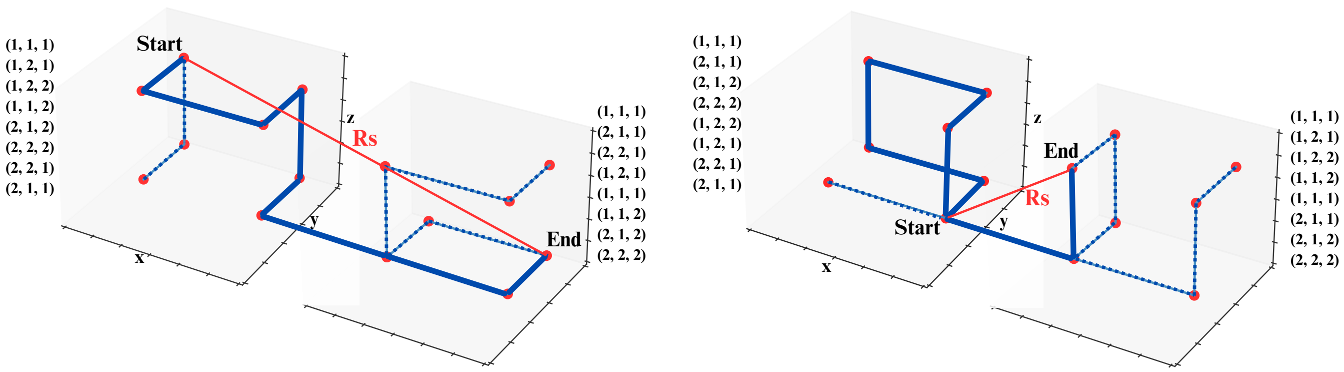


**S1 Figure B.** Upper and lower bounds for end-to-end distances. The left panel shows the connected permissible chromatin conformations in the two adjacent TADs (with 8 bins resolution) which have the largest possible (upper bound) end-to-end distance $R_s(s)$ for the subchain of length $s = 8$. The subchain is designated by 'Start' and 'End' lables. The right panel shows the chromatin conformations with the smallest possible (lower bound) end-to-end distance $R_s(s)$ for the same length $(s = 8)$ of subchain. As in the S1 Figure A, for convenience, the integers are used to label the centers of each bin along the full two-TADs path. We examined all permissible two-TADs conformations with any possible subchain starting point in the first TAD and ending point in the second TAD.

#### 3.2 Excluded distance values

A detailed analysis of the algorithm's output shows that certain values of distance that corresponded to specific pairs of bins never appear in the generated $R_s$ results. One can explain the reason of these exclusions with a lemma from Graph Theory. Imagine the center of each bin as a vertex in a graph, with edges connecting these vertices to symbolize each step in the MC-TAD algorithm (S1 Figure C).

**Lemma 1.** *Assume $G$ is a graph with a set of vertices $V$ and adjacency matrix $A = a_{uv}$. Then the number of uv-walks of the length $k$ in the G(V,E) is the (u, v) entry of $A^k$ [1].*

**S1 Figure C.** Graph representation and adjacency matrix correspond to a cube in the MC-TAD algorithm.

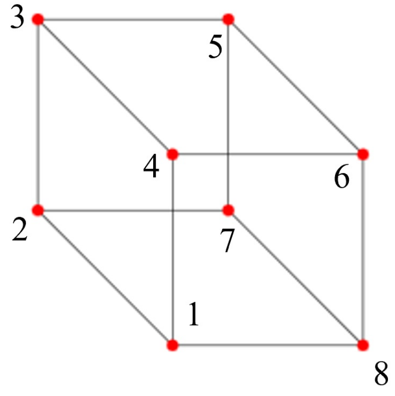


**(a)** Graph representation

$$A = \begin{bmatrix} 0 & 1 & 0 & 1 & 0 & 0 & 0 & 1 \\ 1 & 0 & 1 & 0 & 0 & 0 & 1 & 0 \\ 0 & 1 & 0 & 1 & 1 & 0 & 0 & 0 \\ 1 & 0 & 1 & 0 & 0 & 1 & 0 & 0 \\ 0 & 0 & 1 & 0 & 0 & 0 & 1 & 1 \\ 0 & 0 & 0 & 1 & 1 & 0 & 0 & 1 \\ 0 & 1 & 0 & 0 & 1 & 0 & 0 & 1 \\ 1 & 0 & 0 & 0 & 0 & 1 & 1 & 0 \end{bmatrix}$$

**(b)** Adjacency matrix

**S1 Figure C.** Chromatin conformation at ∼14-kb resolution (8 bins in the cube) can be represented by a graph $G(V, E)$. Vertices correspond to the centers of bins in the cube, which are represented by red circles. All segments are possible and are present or not depending on the geometry of the generated path in the cube. The adjacency matrix $A$ describes the connectivity structure of the graph.

**S1 Figure D.** $A^7$ matrix.

$$A^7 = \begin{bmatrix} 0 & 547 & 0 & 547 & 546 & \underline{0} & \underline{0} & 547 \\ 547 & 0 & 547 & 0 & \underline{0} & 546 & 547 & \underline{0} \\ 0 & 547 & 0 & 547 & 547 & \underline{0} & \underline{0} & 546 \\ 547 & 0 & 547 & 0 & \underline{0} & 547 & 546 & \underline{0} \\ 546 & 0 & 547 & 0 & 0 & 547 & 547 & 0 \\ 0 & 546 & 0 & 547 & 547 & 0 & 0 & 547 \\ 0 & 547 & 0 & 546 & 547 & 0 & 0 & 547 \\ 547 & 0 & 546 & 0 & 0 & 547 & 547 & 0 \end{bmatrix}$$

**S1 Figure D.** The matrix $A^7$ represents the number of paths of length 7 in the graph. The elements in the upper-right block of the matrix correspond to paths that start at one of the left-edge vertices (1, 2, 3, or 4) and end at one of the right-edge vertices (5, 6, 7, or 8) according to S1 Figure C. We highlight these zeros with a red underline to indicate the absence of such paths.

Looking at the elements of the resulting $A^7$ matrix, one can observe in its first 4 rows that the walks, that start from vertex 1, after 7 moves ($k = 7$) will not end in the vertices 6 or 7, see S1 Figure C. Similarly, for starting points at vertices 2, 3, and 4, the path cannot end at certain vertices after 7 moves, according to the zero elements of this matrix.

Since we consider the maximum length of the paths inside the $2 \times 2 \times 2$ cube (number of segments in the corresponding $2 \times 2 \times 2$ graph) to be 7 (maximum length: $2 \times 2 \times 2 - 1$), having zero values in the $\mathrm{A}^7$ matrix indicates that there is no walk with length $k = 7$ between those two ($u$ and $v$) vertices in the graph. For example, if we start from vertex 3 in S1 Figure C, we cannot end up in vertices 1 and 6 and so forth.

Without loss of generality, we only highlighted (with a red underline) the zeros corresponding to the X+ direction to the second cube (vertices 5,6,7 and 8 as ending points). See S1 Figure D.

##### 3.3 Calculating $\sigma_{(2\times2\times2)}$

Upon generating new permissible paths and estimating end-to-end distances $R_i(s)$ (Eq. 2, $s = 8$ case) we use the following convergence criterion to stop the algorithm. Once the algorithm stops to increase the frequencies (number of occurrences) for the set of unique end-end distances, it stops. In the case of paths in two $2 \times 2 \times 2$ cubes, after about 2,000 runs, there is practically no increase in the number of occurrences of the 4 unique end-end distances for the generated conformations (see S1 Figure E and S1 Table A). The distribution of the frequencies for these unique end-to-end distances, and the associated $\sigma$ values, converges even faster, as seen in the S1 Table A.

**S1 Figure E.** Convergence plot for $2 \times 2 \times 2$ cubes.

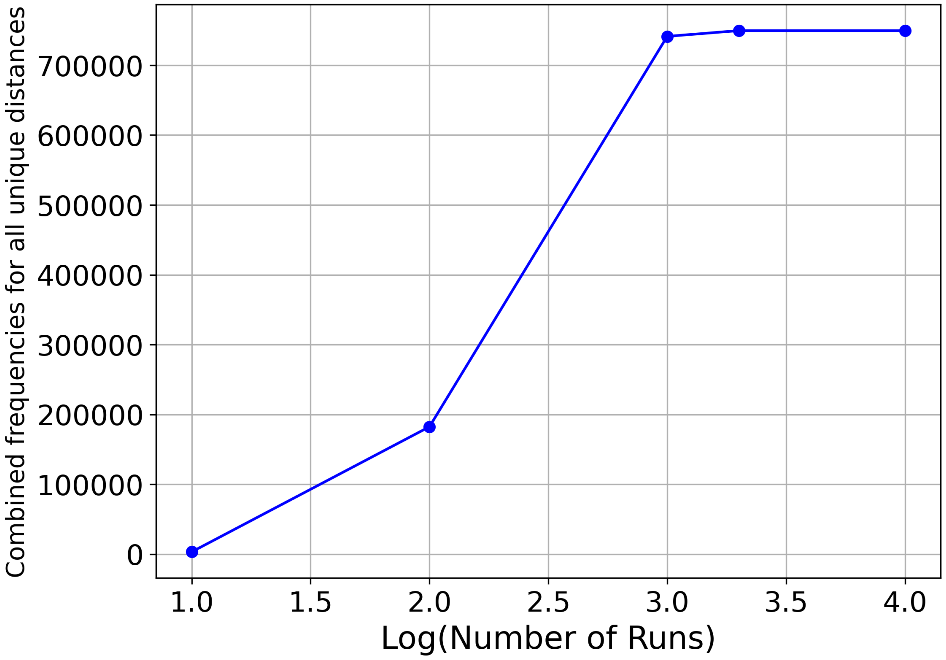


**S1 Figure E.** Convergence of the combined frequencies for all unique end-to-end distances for the paths generated by the MC-TAD algorithm in two adjacent $2 \times 2 \times 2$ cubes. There is no increase in the number of occurrences of unique end-to-end distances after about 2,000 runs.

**S1 Table A.** Distribution of the end-to-end $R_s$ distances in the two $2 \times 2 \times 2$ cubes.

**S1 Table A.** Unique end-to-end distance values and their frequencies (number of occurrences) for the generated paths in two adjacent $2 \times 2 \times 2$ cubes, each of the size of 1.0 arbitrary unit (a.u.). The distribution of the frequencies for these distances is essentially converged after 100 runs.

| Runs | Distance Values (a.u.) | Frequency | $\sigma$ (a.u.) |
|---|---|---|---|
| **100** | 0.707, 1.00, 1.225, 1.58 | 918, 876, 831, 479 | 0.31 |
| **1,000** | 0.707, 1.00, 1.225, 1.58 | 207408, 184466, 211422, 138048 | 0.31 |
| **2,000** | 0.707, 1.00, 1.225, 1.58 | 209792, 186304, 213440, 140032 | 0.31 |

#### 3.4 Calculating $\sigma_{(3\times3\times3)}$ and $\sigma_{(4\times4\times4)}$

Considering paths in the two cubes, the current algorithm formally does not provide convergence at the resolutions higher than 14 kb, corresponding to the 8-bins cube case. The combined number of the frequencies continues to increase with increasing number of runs after $10^3$. However, according to S1 Tables B and C for the $3 \times 3 \times 3$ and $4 \times 4 \times 4$ cubes, the standard deviation ($\sigma$) for the distribution of the frequencies of the unique end-to-end distances for the generated paths remains practically the same across different numbers of runs. This is the consequence of the fact that the ratio of the frequencies for different distinct distance values is preserved across different numbers of runs indicating similar frequency distributions.

**S1 Table B.** Distribution of the end-to-end $R_s$ distances in the two $3 \times 3 \times 3$ cubes.

**S1 Table B.** Unique end-to-end distance values and their frequencies (number of occurrences) for the generated paths in two adjacent $3 \times 3 \times 3$ cubes, each of the size of 1.0 arbitrary unit (a.u.). The distribution of the frequencies for these distances is essentially converged after 100 runs.

| **Runs** | **Distance Values (a.u.)** | **Frequency** | $\sigma$ **(a.u.)** |
|---|---|---|---|
| **100** | 0.33, 0.58, 0.74, 1.0, 1.11, 1.2, 1.37, 1.53, 1.67, 1.73, 1.8, 1.91 | 4014, 6934, 32890, 31948, 26187, 16798, 29968, 13424, 3035, 6048, 3330, 924 | 0.33 |
| **1,000** | 0.33, 0.58, 0.74, 1.0, 1.11, 1.2, 1.37, 1.53, 1.67, 1.73, 1.8, 1.91 | 367476, 628649, 3012267, 2885772, 2440639, 1551073, 2805573, 1249161, 290126, 588601, 299212, 75970 | 0.33 |

**S1 Table C.** Distribution of the end-to-end $R_s$ distances in the two $4 \times 4 \times 4$ cubes.

**S1 Table C.** Distribution of the end-to-end $R_s$ distances in the two $4 \times 4 \times 4$ cubes.
Shown are unique end-to-end distance values and their frequencies (number of occurrences) for the generated paths in two adjacent $4 \times 4 \times 4$ cubes, each of the size of 1.0 arbitrary unit (a.u.). The distribution of the frequencies for these distances is essentially converged after 100 runs.

| **Runs** | **Distance Values (a.u.)** | **Frequency** | $\sigma$ **(a.u.)** |
|---|---|---|---|
| **100** | 0.35, 0.5, 0.61, 0.71, 0.79, 0.87, 0.94, 1.0, 1.06, 1.11, 1.17, 1.22, 1.28, 1.37, 1.46, 1.5, 1.54, 1.58, 1.66, 1.70, 1.77, 1.84, 1.91, 1.97 | 585, 478, 1627, 859, 2633, 376, 2864, 998, 2742, 1702, 736, 779, 3160, 1794, 644, 329, 1206, 576, 250, 426, 407, 450, 91, 80 | 0.34 |
| **1,000** | 0.35, 0.5, 0.61, 0.71, 0.79, 0.87, 0.94, 1.0, 1.06, 1.11, 1.17, 1.22, 1.28, 1.37, 1.46, 1.5, 1.54, 1.58, 1.66, 1.70, 1.77, 1.84, 1.91, 1.97 | 60806, 51527, 183873, 98149, 280270, 45840, 329171, 104190, 321715, 196655, 74945, 93217, 370800, 237606, 72959, 45609, 168229, 87700, 41359, 50599, 52511, 56084, 11670, 10852 | 0.34 |

#### Extension to higher resolution

The maximum theoretically possible, but still biologically meaningful, genomic resolution is a single base-pair, which corresponds to dividing the 100 kb TAD into 100,000 bins (a cube with $46 \times 46 \times 46$ bins). Here we provide a rough estimate for the value of $\sigma_{(46\times46\times46)}$. Obviously, as the number of bins (resolution) is increased, larger end-to-end distances become possible, increasing their spread and the corresponding $\sigma$ values. This leads us to the following reasonable assumption:

$$\sigma_{(2\times2\times2)} < \sigma_{(3\times3\times3)} < \ldots < \sigma_{(46\times46\times46)} \,. \tag{4}$$

That is, we assume that $\sigma_{(\mathrm{N}\times\mathrm{N}\times\mathrm{N})}$ is a monotonic function of $N$. Indeed, we have seen already that $\sigma$ values, $\sigma_{(2\times2\times2)} = 0.31$, $\sigma_{(3\times3\times3)} = 0.33$ and $\sigma_{(4\times4\times4)} = 0.34$.

To estimate $\sigma_{(46\times46\times46)}$, we introduce the maximum, $d_{\max(2\times2\times2)}$, and the minimum, $d_{\min(2\times2\times2)}$, end-to-end distances of the paths in the two cubes with 8 bins ($N = 2$) each. The specific values of these quantities are $d_{\max(2\times2\times2)} = 1.58$ and $d_{\min(2\times2\times2)} = 0.71$, see S1 Table A. The difference between these two values provides the range for possible distances:

$$\mathrm{Range}_{(2\times2\times2)} = d_{\max(2\times2\times2)} - d_{\min(2\times2\times2)} = 0.87 \,. \tag{5}$$

Obviously, $\sigma_{(\mathrm{N}\times\mathrm{N}\times\mathrm{N})} < \mathrm{Range}_{(\mathrm{N}\times\mathrm{N}\times\mathrm{N})}$. But let us make a stronger assumption that these two quantities ($\sigma$ and Range) are proportional to each other. Then

$$\sigma_{(46\times46\times46)} = \left( \frac{\sigma_{(2\times2\times2)}}{\mathrm{Range}_{(2\times2\times2)}} \right) \times \mathrm{Range}_{(46\times46\times46)} \,. \tag{6}$$

Providing that $\mathrm{Range}_{(46\times46\times46)} \approx 2.45$ (the largest distance between the two corners of two adjacent cubes of unit size), the above equation gives us the following rough estimate: $\sigma_{(46\times46\times46)} = 0.87$.

These estimates of $\sigma$ (0.31, 0.33, 0.34, ..., 0.87) demonstrate that while its values increase with increasing model resolution, there is an upper bound to it: 0.87. All $\sigma$ and

Range values discussed here are presented in arbitrary units (1 a.u. = cube size). In the next section, we convert these values into units of $\mu$m.

#### Conversion of arbitrary units of length to micrometers within MC-TAD algorithm

In the previous sections, the side of the cube $l$ was set to $l = 1$ a.u., resulting in the value for the standard deviation $\sigma_{(2\times2\times2)} = 0.31$ a.u. To convert a.u. to $\mu$m, we equate the volume of the average TAD, $V_{\mathrm{TAD}} = \frac{4}{3}\pi R_{\mathrm{TAD}}^3$, to the volume of the cube, $V_{\mathrm{cube}} = (C_{\mathrm{conv}} l)^3$, where $C_{\mathrm{conv}}$ is the conversion factor that can be expressed as:

$$C_{\mathrm{conv}} = \sqrt[3]{\frac{\frac{4}{3}\pi R_{\mathrm{TAD}}^3}{l^3}} = 1.61 \times \left(\frac{R_{\mathrm{TAD}}}{l}\right) . \tag{7}$$

Here and in what follows, we use uppercase $\Sigma$ for standard deviation measured in $\mu$m, to differentiate it from the standard deviation in a.u., denoted by lowercase $\sigma$, see Sec. “Calculating the width $\sigma$ for the distribution of allowed conformations" in “Methods" of the main text. Providing that $l = 1$ a.u., we can convert the standard deviation $\sigma$ in a.u. to the standard deviation $\Sigma$ in $\mu$m using the following relation:

$$\Sigma_{(\mathrm{s=118kb})(\mathrm{14kb})} = 1.61 \times R_{\mathrm{TAD}} \times \sigma_{(2\times2\times2)} . \tag{8}$$

Here, $\Sigma$ carries two indices: the genomic distance ($s = 118$ kb) for which $\sigma_{(2\times2\times2)}$ was calculated, and which corresponds to the average TAD size in X chromosome, and the genomic resolution (14 kb) that corresponds to the size of the bin in the 8-bin $(2 \times 2 \times 2)$ cube (genomic length $s = 1$ in the bin).

Provided that the average radius of a TAD ($R_{\mathrm{TAD}}$) in X chromosome is 0.07 $\mu$m for Li et al., 2017 models and 0.09 $\mu$m for Tolokh et al., 2023 models, one can obtain the following estimates for $\Sigma_{(\mathrm{s=118kb})(\mathrm{14kb})}$ values for these two sets of models: 0.03 $\mu$m and 0.04 $\mu$m, respectively.

In the case of higher resolutions, $3 \times 3 \times 3$ or $4 \times 4 \times 4$, the resolution index values are 4 kb and 2 kb, respectively. For the highest experimentally achievable resolutions 1-2 kb [2], having $\sigma_{(4\times4\times4)} = 0.34$, one can get the following estimate for $\Sigma$ in the case of Tolokh et al., 2023 models (with $R_{\mathrm{TAD}} = 0.09$ $\mu$m):

$$\Sigma_{(\mathrm{s=118kb})(\mathrm{2kb})} = 1.61 \times R_{\mathrm{TAD}} \times \sigma_{(4\times4\times4)} = 0.05\,\mu\mathrm{m}\,. \tag{9}$$

Similar estimation of $\Sigma$ can be obtained in the case, when the cube (the average TAD in X chromosome) is divided into approximately 100,000 bins ($\approx 46^3$), resulting in 1 base pair bin size (resolution). Using our estimated value of $\sigma_{(46\times46\times46)} = 0.87$, and the average TAD radius 0.09 $\mu$m (for Tolokh et al., 2023 models), we arrive at:

$$\Sigma_{(\mathrm{s=118kb})(\mathrm{1b})} = 1.61 \times R_{\mathrm{TAD}} \times \sigma_{(46\times46\times46)} = 0.13\,\mu\mathrm{m}\,. \tag{10}$$

#### Conversion of arbitrary units of length to micrometers for Ulianov et al., 2021 models

In Ulianov et al., 2021 models the arbitrary units (a.u.) of length are used, each corresponding to the size of bead which models a 10 kb portion of chromatin in a TAD. To convert these a.u. to $\mu$m, we assume that the 10 kb portion of chromatin corresponds to a chain of 50 nucleosomes (each containing 200 bp of DNA) separated (center-to-center) by 15 nm [3]. In addition, we assume that the conformation of a nucleosome chain at a small scale inside interphase TAD follows a random walk conformation [4]. This leads us to the following estimate for the size (diameter) of 10 kb (50 nucleosome) bead in Ulianov et al., 2021 models of X chromosome:

$$1\,\mathrm{a.u.} = \sqrt{50} \times 15\,\mathrm{nm} = 106.06\,\mathrm{nm} = 0.106\,\mu\mathrm{m}\,. \tag{11}$$

#### Relative *Conformational Heterogeneity* for the three models of fruit fly X chromosome

**S1 Table D.** *Relative C.H.* values for the three models of X chromosome.

**S1 Table D.** *Relative Conformational Heterogeneity* values at different genomic separations for the three models of X chromosome at approximately 10-kb resolution, see Fig 9b in the main text. The error bars represent the standard deviations estimated using 10 sets of chromatin conformations for the Li et al., 2017 and Tolokh et al., 2023 models. Genomic separation $s = 118$ kb corresponds to the average genomic size of TAD in X chromosome.

| **Models of X chromosome** | **14 kb** | **118 kb** | **1 Mb** | **10 Mb** |
|---|---|---|---|---|
| Tolokh et al., 2023 | 0 | $0.14 \pm 0.01$ | $0.05 \pm 0.02$ | $0.19 \pm 0.04$ |
| Li et al., 2017 | 0 | $0.15 \pm 0.01$ | $0.06 \pm 0.02$ | $0.24 \pm 0.07$ |
| Ulianov et al., 2021 | 0.04 | 0.15 | 0.16 | 0.08 |

#### Estimating *Relative C.H.* at 1-bp resolution

**S1 Figure F.** *Relative* $\langle R_s \rangle$ at 1-bp resolution.

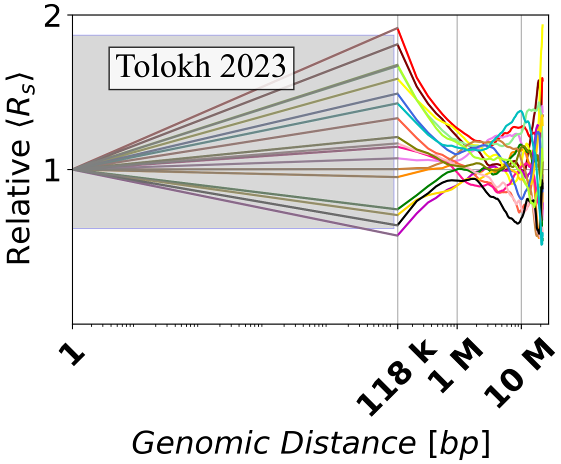


**S1 Figure F.** Extrapolation of the ensemble X chromosome conformations from Tolokh et al., 2023 models to the theoretical limit of 1-bp resolution, corresponding to $\Sigma_{(\mathrm{s=118kb})(\mathrm{1bp})} = 0.13\ \mu$m. Shown are the dependencies of the *relative* mean Euclidean inter-particle distances (*Relative* $\langle R_s \rangle$) on the genomic separation $s$ between the polymer beads calculated using Eq 4 of the main text. In the shaded region a linear interpolation is used to guide the eye, see “Methods" in the main text.

**S1 Table E.** *Relative C.H.* for Tolokh et al., 2023 chromatin models [5] at 1-bp resolution.

**S1 Table E.** Values of the *Relative C.H.* at 1-bp resolution (with the corresponding value of $\Sigma_{(\mathrm{s=118kb})(\mathrm{1bp})} = 0.13\,\mu\mathrm{m}$) estimated for several genomic separations for Tolokh et al., 2023 chromatin models [5]. 118 kb separation represents the average size of TAD in the X chromosome, at which the spread of $\langle R_s \rangle$ values for different chromosome conformations is calculated. See "Methods". The error bars are the standard deviations over all 10 values obtained as the *Relative C.H.* of different sets of selected chromatin ensembles.

| **Models of X chromosome** | **1 bp** | **118 kb** | **1 Mb** | **10 Mb** |
|---|---|---|---|---|
| Tolokh et al., 2023 | 0 | $0.41 \pm 0.01$ | $0.11 \pm 0.020$ | $0.2 \pm 0.04$ |

#### Comparing the characteristic size of fruit fly X chromosome in the three models

The relatively low *Relative C.H.* at the largest (20 MB) genomic distances in Ulianov et al., 2021 model, compared to that of the Li et al., 2017 and Tolokh et al., 2023 models, can be explained by the distinctly smaller size of the volume available to the X chromosome in Ulianov et al., 2021 models compared to the other two models (see the explanation in the main text, S1 Figure G and S1 Table F in the S1 text below). As noted in the main text, we suggest that the smaller chromosome size in Ulianov et al., 2021 models is a consequence of a smaller simulation box used, consistent with the maximum extensions seen the S1 Table F. Note that confinement (box size) becomes important when the natural, that is unconstrained, polymer size is larger or comparable to the dimensions of the confining box, which is the case for all three models. Comparing the box sizes across models, the Ulianov et al., 2021 models use a 22 DPD a.u. box (2.2 $\mu$m, $V = 10.65\,\mu\text{m}^3$), while the Tolokh et al., 2023 models have a $4 \pm 0.12\,(\mu\text{m})$ sphere size ($33.5 \pm 3\ \mu\text{m}^3$), making the confining box volume for Ulianov et al., 2021 models more than three times smaller. The relatively limited space in the Ulianov et al., 2021 models restricts chromatin movement, leading to less divergence of the polymer chain configurations, which explains the low *Relative C.H.* at large ($10 - 20$ Mb) genomic distances.

**S1 Figure G.** Average end-to-end distances of X chromosome across the three models.

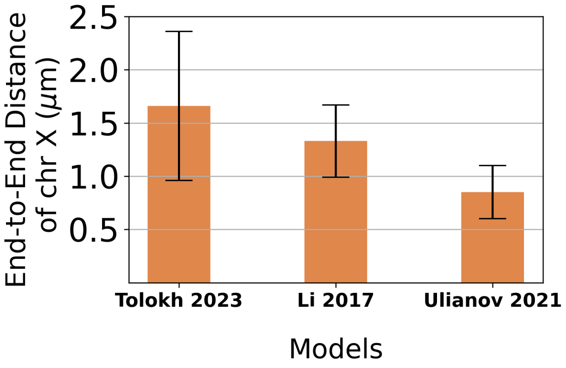


**S1 Figure G.** Average end-to-end distances of X chromosome in the three different models. (Heterochromatin and centromere regions are not included in all three models). Error bars represent standard deviations for one ensemble selection (∼20 chromatin conformations) in the models of Tolokh et al., 2023 and Li et al., 2017, and the standard deviation for a single ensemble of 20 nuclei in the model of Ulianov et al., 2021. See Sec. "Chromosome Size" in "Methods".

**S1 Table F.** Maximum extension and average end-to-end distance of the Drosophila X chromosome across the three models.

**S1 Table F.** Maximum extension and average end-to-end distance of the Drosophila X chromosome in the three models. These values are averaged over ∼20 single cells, with error bars representing the standard deviation across the ensemble.

| **Models of X chromosome** | **Max Extension (micrometers)** | **End-to-End Distance (micrometers)** |
|---|---|---|
| Ulianov et al., 2021 | 1.53 | $0.85 \pm 0.25$ |
| Tolokh et al., 2023 | 4.11 | $1.66 \pm 0.7$ |
| Li et al., 2017 | 3.85 | $1.33 \pm 0.34$ |

#### 3.5 A simplified example demonstrating non-uniqueness of correspondence between bulk and single-cell Hi-C matrices

For example, consider the matrix $A$ and three "single-cell" binary matrices $S_1$, $S_2$ and $S_3$:

$$A = \begin{pmatrix} 1 & 0.45 & 0 \\ 0.45 & 1 & 0 \\ 0 & 0 & 1 \end{pmatrix}, \quad S_1 = \begin{pmatrix} 1 & 1 & 0 \\ 1 & 1 & 0 \\ 0 & 0 & 1 \end{pmatrix}, \quad S_2 = \begin{pmatrix} 1 & 0 & 0 \\ 0 & 1 & 0 \\ 0 & 0 & 1 \end{pmatrix}, \quad S_3 = \begin{pmatrix} 1 & 0.5 & 0 \\ 0.5 & 1 & 0 \\ 0 & 0 & 1 \end{pmatrix} \tag{12}$$

There is more than one set of $p_1$, $p_2$ and $p_3$ such that:

$$\begin{pmatrix} 1 & 0.45 & 0 \\ 0.45 & 1 & 0 \\ 0 & 0 & 1 \end{pmatrix} = p_1 \begin{pmatrix} 1 & 1 & 0 \\ 1 & 1 & 0 \\ 0 & 0 & 1 \end{pmatrix} + p_2 \begin{pmatrix} 1 & 0 & 0 \\ 0 & 1 & 0 \\ 0 & 0 & 1 \end{pmatrix} + p_3 \begin{pmatrix} 1 & 0.5 & 0 \\ 0.5 & 1 & 0 \\ 0 & 0 & 1 \end{pmatrix}.$$

For example:

1. $p_1 = 0.3$, $p_2 = 0.4$, $p_3 = 0.3$
2. $p_1 = 0.4498$, $p_2 = 0.5498$, $p_3 = 0.0004$
